# Brain and Organoid Manifold Alignment (BOMA), a machine learning framework for comparative gene expression analysis across brains and organoids

**DOI:** 10.1101/2022.06.13.495946

**Authors:** Chenfeng He, Noah Cohen Kalafut, Soraya O. Sandoval, Ryan Risgaard, Chen Yang, Saniya Khullar, Marin Suzuki, Qiang Chang, Xinyu Zhao, Andre M.M. Sousa, Daifeng Wang

## Abstract

Organoids have become valuable models for understanding cellular and molecular mechanisms in human development including brains. However, whether developmental gene expression programs are preserved between human organoids and brains, especially in specific cell types, remains unclear. Importantly, there is a lack of effective computational approaches for comparative data analyses between organoids and developing humans. To address this, by considering the public data availability and research significance, we developed a machine learning framework, Brain and Organoid Manifold Alignment (BOMA) for comparative gene expression analysis of brains and organoids, to identify conserved and specific developmental trajectories as well as developmentally expressed genes and functions, especially at cellular resolution. BOMA first performs a global alignment and then uses manifold learning to locally refine the alignment, revealing conserved developmental trajectories between brains and organoids. Using BOMA, we found that human cortical organoids better align with certain brain cortical regions than other non-cortical regions, implying organoid-preserved developmental gene expression programs specific to brain regions. Additionally, our alignment of non-human primate and human brains reveals highly conserved gene expression around birth. Also, we integrated and analyzed developmental scRNA-seq data of human brains and organoids, showing conserved and specific cell trajectories and clusters. Further identification of expressed genes of such clusters and enrichment analyses reveal brain- or organoid-specific developmental functions and pathways. Finally, we experimentally validated important specific expressed genes using immunofluorescence. BOMA is open-source available as a web tool for general community use.

## Introduction

The development of human brains, especially during the early periods, remains poorly understood^1–3^. Understanding how neural stem cells differentiate into the myriad cell types that form the brain, especially at the molecular level, such as gene expression and its regulatory mechanisms, will shed light on the human brain development and potentially further help understand the etiology of neurodevelopmental diseases. Several large collaborative consortia have been carried out to generate large-scale next generation sequencing data in human brains, aiming to provide functional genomic resources for understanding molecular mechanisms of human brain and brain development. For example, BrainSpan^4^ collected ∼600 tissue samples from 48 postmortem human brains, ranging from prenatal to adult age groups and measured the transcriptomic and epigenomic data across developmental stages and brain regions. PsychENCODE^1,5^ generated multi-omics data for approximately 2,000 postmortem brains, aiming to understand functional genomics and gene regulation in human adult brains and neuropsychiatric disorders. These consortia provided valuable public resources to decipher the developmental functional genomics and gene regulation in the human brain. However, the postmortem brain samples serve only as snapshots of the brain at different stages, whereas brain development is a dynamic process that requires crosstalk among various genes, cell types, brain regions and environment^6^.

Because it is quite challenging to measure *in-vivo* molecular activities such as gene expression in human brains, animals such as rodents and non-human primates (NHP) have been used as models for studying molecular mechanisms during brain development. For example, Zhu et al.^7^ have thoroughly compared the bulk RNA-seq data of brain development between humans and rhesus macaques at multiple brain regions and time points. Particularly, they performed Non-negative Matrix Factorization to linearly factorize the gene expression matrix into five biologically meaningful ‘transcriptomic signatures’, which were then compared between human and NHP. Such comparisons demonstrated the usefulness of NHP models for studying brain development. However, their results also highlighted the divergence of molecular mechanisms across species. This is also supported by previous studies that noticed that using animals as models is insufficient because brain maturation is specific to its developmental context^4,8^ and human brains have specific developmental programs that allow, for example, a dramatic size expansion when compared to other primates^9^.

To solve these challenges, emerging 3D brain culture technologies, such as organoids, have been developed. These cultures utilize embryonic (ESCs) or induced Pluripotent Stem Cells (iPSC) and differentiate them into 3D human brain models^10^. An intriguing discovery is that iPSCs follow intrinsic programs and extrinsic cues to form 3-dimensional forebrain organoids (3DOs) that can be maintained for at least 40 weeks and even over two years, with a transcriptomic signature corresponding to “birth” at ∼28 weeks of culture^11,12^. Organoids as brain models, although in their early developing stages, have already found numerous medical applications. For example, Park et al. used 1,300 organoids to model the human brain and conducted drug screening for Alzheimer’s disease^13^. However, to what extent the *in-vitro* cultured organoids preserve the *in-vivo* complex dynamic process remains a question^14^, with contradictory conclusions that have been made by the community. For example, Gordon et al.^11^ cultured organoids for up to 694 days and used Transition Mapping (TMAP), a rank-rank hypergeometric test based method, to map the organoids bulk RNA-seq datasets with BrainSpan RNA-seq datasets and demonstrated organoids culture could reproduce several developmental milestones of *in-vivo* brain development even at mid-to late-fetal stages. Velasco et al.^15^ performed single-cell RNA-seq (scRNA-seq) on 166,242 cells isolated from 21 organoids and showed that organoids can virtually indistinguishably reproduce the cell diversity of the human cerebral cortex. On the other hand, Pollen et al.^9^ compared human primary tissues versus human organoids, using canonical correlation analysis (CCA)^16^ and co-clustering of the mixture cells from both origins. They found organoids maintained the composition of cell types but varied in the cell percentages and concluded that using organoids as brain models is promising, but the organoids protocols need future improvements to better preserve the brain cell type fractions and cell functions^9,17^. Bhaduri et al.^18^ compared single-cell gene expression data of samples across different developmental periods and multiple cortical areas with organoids and found cellular stress pathways have been activated in organoids, which impairs cell-type specification during organoid development. All these studies highlight the promise of using organoids as models for brain developmental research; however, until now, the fidelity of organoid models is still under debate. One of the attributed reasons is the lack of dedicated computational approaches for integrative and comparative analysis of gene expression across developmental stages between brains and organoids, especially for single-cell datasets.

In particular, the comparative analysis of developmental data between brains and organoids can be viewed by machine learning as an alignment problem across multiple datasets. For instance, manifold alignment^19,20^, a popular machine learning technique, projects samples from multiple datasets onto a common latent space via mapping manifolds across datasets, e.g., multi-omics datasets^21^. The neighboring samples on the latent space suggest that they can be aligned and thus share similar features (or distant samples for unaligned). In general, manifold alignment algorithms^22^ can be supervised or unsupervised depending on whether the sample correspondence is provided (supervised) or not (unsupervised). A supervised approach needs predefined correspondence between the samples across two datasets^23^. For example, ManiNetCluster^24^ embeds samples into a latent manifold space and aligns them by minimizing the overall distances between corresponding samples. An unsupervised approach does not require correspondence, instead learn the correspondence across multiple datasets^25^. For example, MATCHER^21^ performs linear trajectory alignment based on latent Gaussian process; MMD-MA^26^ maximizes mean discrepancy on a kernel space. UnionCom^27^ uses matrix optimization to match the distance matrices of each dataset. SCOT^28^ incorporates Gromov Wasserstein-based optimal transport to align single-cell datasets. However, the unsupervised approaches, in general, automatically assume a shared underlying structure among the aligned datasets^29^, which might not always be true. Besides, none of these manifold alignment methods considered prior time information across samples in development that can likely help increase performance and interpretability^30^ of the alignment. The developmental data for brains or organoids typically provide prior time information on developmental stages, e.g., Postconceptional weeks (PCWs) of developing brains, and cultured days of organoids. Such prior time information, though at low resolution, may help predict initial correspondences globally across samples from different datasets, in contrast to the fully unsupervised fashion. Building on such initial correspondence, further manifold alignment can then refine the alignment to reveal higher resolution and local timing by the manifold shapes that have been widely used to uncover pseudo timings^31^. However, to the best of our knowledge, manifold alignment has yet been applied for integrative and comparative analysis of brain and organoid data, especially for development and single cells.

In this paper, we developed a manifold learning framework, Brain and Organoid Manifold Alignment (BOMA), to align developmental gene expression data across human brains and organoids, aiming to better understand conserved and specific gene expression and functions. In particular, BOMA adopts a semi-supervised manifold alignment manner. That is, using prior timing information from datasets, we first perform a global alignment at a coarse-grained level to generate a correspondence matrix among samples. Next, a non-linear manifold alignment is performed to refine the alignment and identify the developmental trajectories comprising cross-dataset samples with higher resolution, local pseudo times. The aligned and unaligned samples aim to uncover conserved and specific developmental trajectories across human brains and organoids. We first demonstrated an application of BOMA by aligning bulk RNA-seq gene expression datasets and observed a similar developing trend as in the original respective publications^11^. By aligning organoids with different human brain regions, we also found that organoids are more similar to certain brain regions at specific time points. We also aligned the scRNA-seq data of human versus chimpanzee organoids and observed a delayed development of human organoids compared with chimpanzee organoids. Finally, we compared recent time-series scRNA-seq datasets between human brains and human organoids in development. We found both common and uniquely expressed genes between the brains and organoids at resolutions of cell types across developmental stages. Moreover, we experimentally validated the expression of genes displaying differences between brains and organoids in selected cell types. BOMA is also available as an open-source web tool for community use.

## Results

### Brain-Organoid Manifold Alignment (BOMA) framework for comparative analyses of gene expression data between brains and organoids

As shown in Figure 1, BOMA inputs developmental gene expression matrices of the brain and organoid samples (e.g., tissues, cells). First, it uses global alignment to align the samples and initialize a sample-wise correspondence matrix at a coarse-grain level (e.g., via manifold warping). Second, BOMA performs a manifold alignment using the correspondence matrix as the initial alignment. This step finds shared manifolds of the samples and maps them onto a common manifold space. The manifold shapes of the samples on the space are expected to uncover various developmental trajectories, which can be either conserved across brains and organoids (aligned samples) or brain/organoid specific (unaligned samples). We also designed an alignment score, which is derived from the Euclidean distances of samples on the common space, to quantify the similarities across samples. Finally, BOMA clusters the samples on those trajectories and alignment scores and finds underlying differentially expressed genes (DEGs), enriched gene functions, and associated phenotypes for each cluster, providing a deeper understanding of developmental functional genomics in brains vs. organoids. The full description of the BOMA model is available in Methods and Materials.

**Figure 1.**
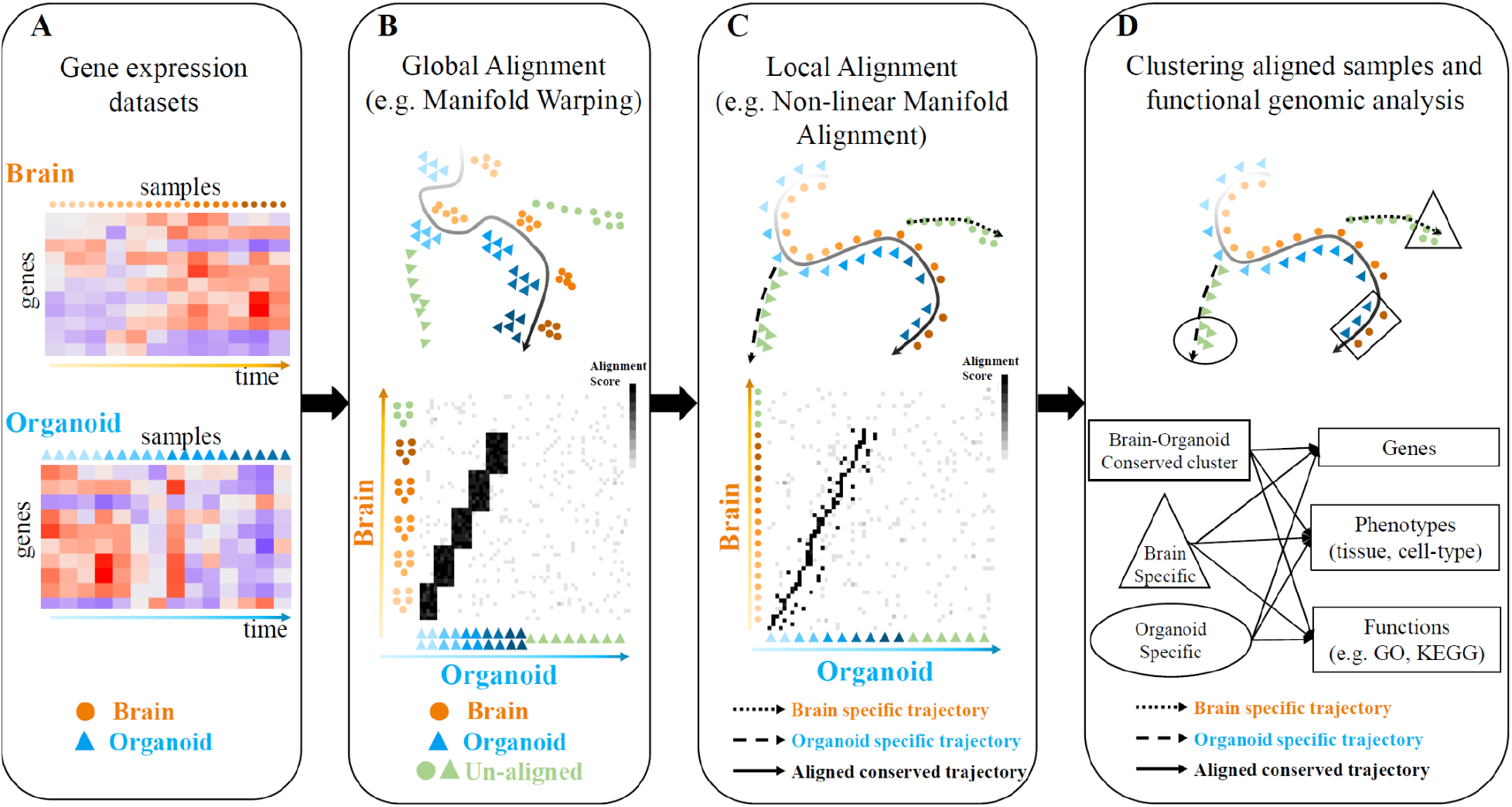
Brain and Organoid Manifold Alignment (BOMA), a computational framework for comparative analyses of gene expression data between brains and organoids. (**A**) BOMA inputs multiple gene expression datasets (genes by samples) from brains and organoids. The samples are ordered by prior timing information. **(B)** Step 1 - Global alignment to infer the correspondences of samples across the datasets at a coarse-grain level. **(C)** Step 2 – Local alignment to refine the alignment and map samples onto a common manifold space. (**D**) Clustering and functional analysis of aligned samples on the common space, e.g., brain-organoid conserved (square) or specific (circle and triangle) clusters and developmental trajectories (black curves). Downstream analyses of those clusters can discover differentially expressed genes, enriched gene functions, and associated phenotypes.

### Spatiotemporal conservation and divergence of gene expression between organoid and brain regions

Recent landmark studies compared gene expression between the human brain and organoid development^11^. However, our understanding of where and when gene expression in various brain regions is conserved or different from organoids is still unclear. To this end, we applied BOMA to align developmental gene expression data of human brains and organoids at the bulk tissue level. The brain dataset includes brain tissue samples in BrainSpan^4^ (N=460, Dataset 1, Table S1) from 16 human brain regions (Table S2). The organoid dataset includes organoids from a recently published long-term cultured ‘human cortical spheroid (hCS)’ organoid bulk RNA-seq dataset^11^ (N=62, Dataset 6, Table S1).

Our alignment shows these brain and organoid samples primarily follow a shared trajectory on the common space, indicating potential conservation during their development (Figure 2A). In particular, as shown in Figure 2B, the organoid samples from 25 days to 250 days were aligned with the brain tissue samples at prenatal stages^11^. At 300 days, the organoid samples started to gain postnatal signatures, indicated by their high alignment scores with postnatal brain samples (Figure 2B). Also, organoids after 350 days were not well aligned with any brain samples, which indicates that the late-stage organoids may differ from postnatal brain development. This observation was consistent with a recent comparison of brains and organoids development^11^.

**Figure 2.**
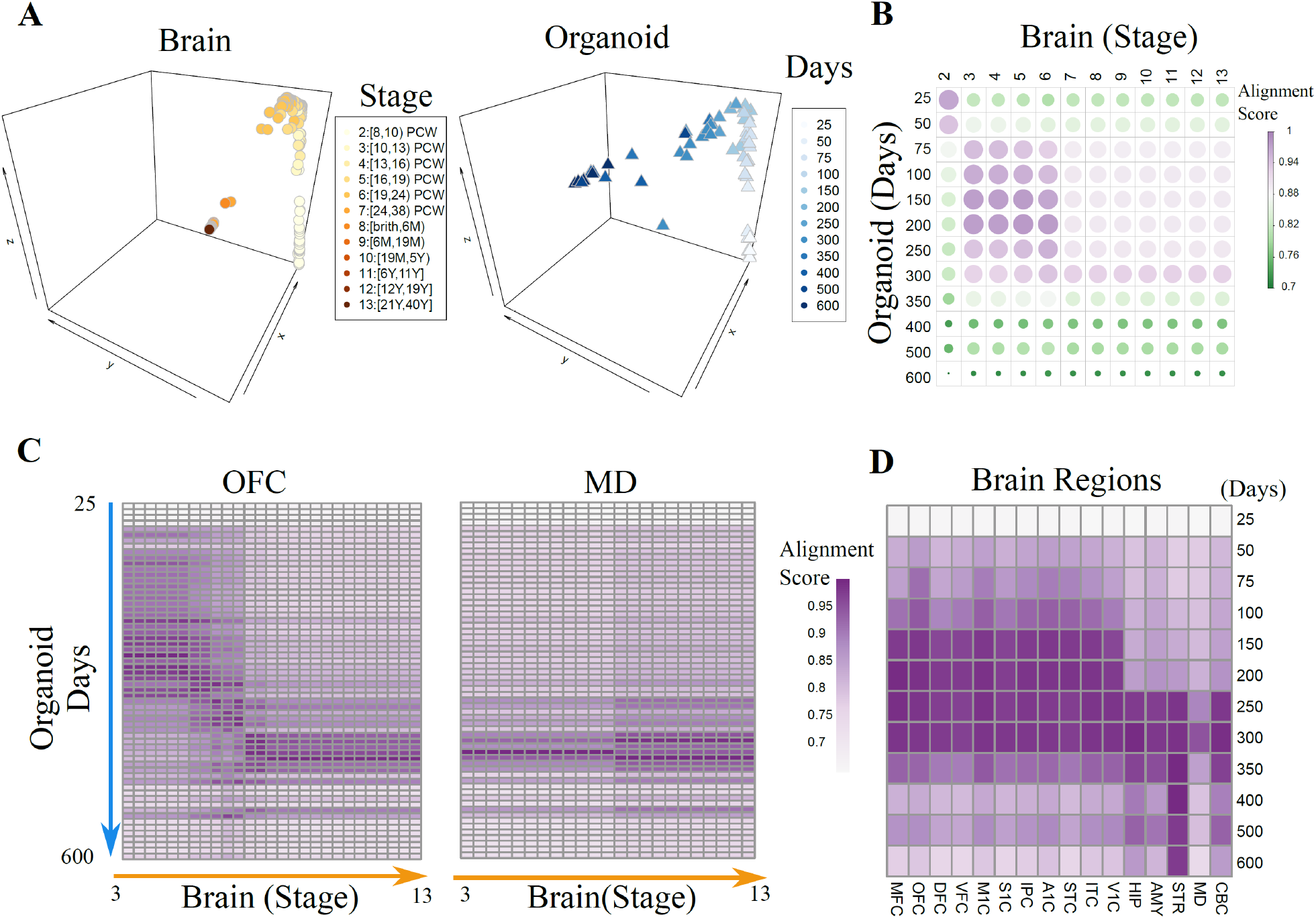
Spatiotemporal conservation and divergence of gene expression between organoid and brain regions. (**A**) Aligned human brain^4^ and organoid^11^ samples on the common space from BOMA. Human brain samples are from BrainSpan and are colored in orange by developmental stages as described by Kang et al.^32^. Organoid samples are colored in blue by the cultured days. PCW: Post Conception Weeks; M: Month; Y: Year. (**B**) Correlation plot shows the similarity (quantified as ‘Alignment Score’, Methods) of aligned samples between the brain and organoid samples. Each dot is the averaged similarity across all pairs of samples at the specific developmental time points. (**C**) Alignment scores between organoids with brain samples from Orbital frontal cortex (OFC) and Mediodorsal thalamus (MD). (**D**) Averaged alignment scores between organoids versus the 16 brain regions in BrainSpan.

Furthermore, we also interrogated which human brain regions are most similar to organoids in development. To answer this, we assessed the BOMA alignment of brain samples from each individual region with the organoid samples. As expected, distinct alignment patterns were found for different brain regions, with cortical areas aligning better with hCS organoids than non-cortical brain regions during early developmental stages up to 200 days (Figure 2C). At 200 days, the alignment scores of cortical regions are significantly higher than non-cortical regions (one-side t-test p<6.9e-4). However, within the neocortex, several cortical areas like Orbital Frontal Cortex (OFC), Inferior Parietal Cortex (IPC), Primary Auditory Cortex (A1C), Superior Temporal Cortex (STC) are more aligned with cortical organoids than other cortical areas up to 100 days (one-side t-test p<1.5e-4; Figure 2D, Figure S1). It is important to note that because several brain regions did not have samples from Stage 2, we removed Stage 2 from all the regions for the BOMA alignment. As a result, organoids at 25 days cannot align with any brain samples.

Therefore, these results suggest that cortical organoids specifically preserve brain-regional development at certain stages (i.e., spatiotemporal conservation), instead of mimicking the whole brain development. Interestingly, they also suggest that organoids are transcriptomically closer to certain neocortical areas, particularly perisylvian and orbital frontal areas, than they are to other neocortical areas.

### Developmental gene expression discrepancies between human and chimpanzee organoids

We also applied BOMA to align human brains and Non-Human Primates (NHP) brains, revealing their conserved developmental gene expression across species. Specifically, we aligned rhesus macaque brain samples (N=366)^7^ with human brain samples (N=460) from BrainSpan using BOMA. We found those samples were aligned most closely around the time of birth (Figure S2), indicating that brain gene expression across two species becomes more similar at the bulk tissue level perinatally^7^.

To deepen our understanding of the conservation and specificity of developmental gene expression at cellular resolution across human and NHP organoids, we applied BOMA to developmental scRNA-seq data and aligned single cells of human organoids (N=47,130, Dataset 7, Table S1) versus chimpanzee organoids (N=26,228, six time points, Dataset 7, Table S1)^33^. The BOMA alignment was performed on the pseudo-cells identified by the same approach as the study generated the datasets^33^, aiming to combat single-cell expression noises. Each pseudo-cell represents a group of cells with similar gene expression patterns. In total, 938 human and 483 chimpanzee organoid pseudo-cells were generated for BOMA. Our analysis shows that these pseudo-cells from the two species organoids were aligned in general to a common trajectory, which indicates cross-species developmental similarity in organoids (Figure 3A). However, some discrepancies could also be observed. First, compared with human cells, chimpanzee cells were shifted towards a later time over the maturational trajectory (towards the left in Figure 3A), suggesting that chimpanzee organoids were developing faster than human organoids (Figure 3B). The observed protracted maturation of human organoids is in line with the previous study^33^, and was also observed in other cross-species comparison studies on organoids^34,35^ as well as on 2D cultures^36,37^. Second, we noticed that two sets of chimpanzee cells (Chimpanzee_1 and Chimpanzee_2 in the right panel of Figure 3A) could not be well aligned with any human cells. To understand the functional relevance of these two sets of cells (Data 1), we extracted upregulated genes and performed a functional enrichment analysis (Methods, Figure 3C). Genes upregulated in Chimpanzee_2 were mostly enriched with brain developmental functions (FDR < 10e-5). For example, the most significantly enriched term, ‘Neuron projection morphogenesis’ is related to the maturation of neurons and circuit assembly^38^. These observations again indicate a faster maturation of chimpanzee organoids. On the other hand, upregulated genes in Chimpanzee_1 at early time points (0 and 4 days) were enriched in cell division processes (e.g., cell cycle, chromatin remodeling, etc.). This suggests cell division and proliferation are dramatically more active in chimpanzee than in human at early developmental stages *in vitro*, which agrees with the previous finding that metaphase in human cerebral cortex progenitor cells is significantly longer than in chimpanzees^35^. Since progenitor cells in human have longer cell division, their differentiation into postmitotic neurons is slower^33^. Therefore, the delayed development of human organoid cells in Chimpanzee_2 can be traced back to the functional distinction of cells in Chimpanzee_1.

**Figure 3.**
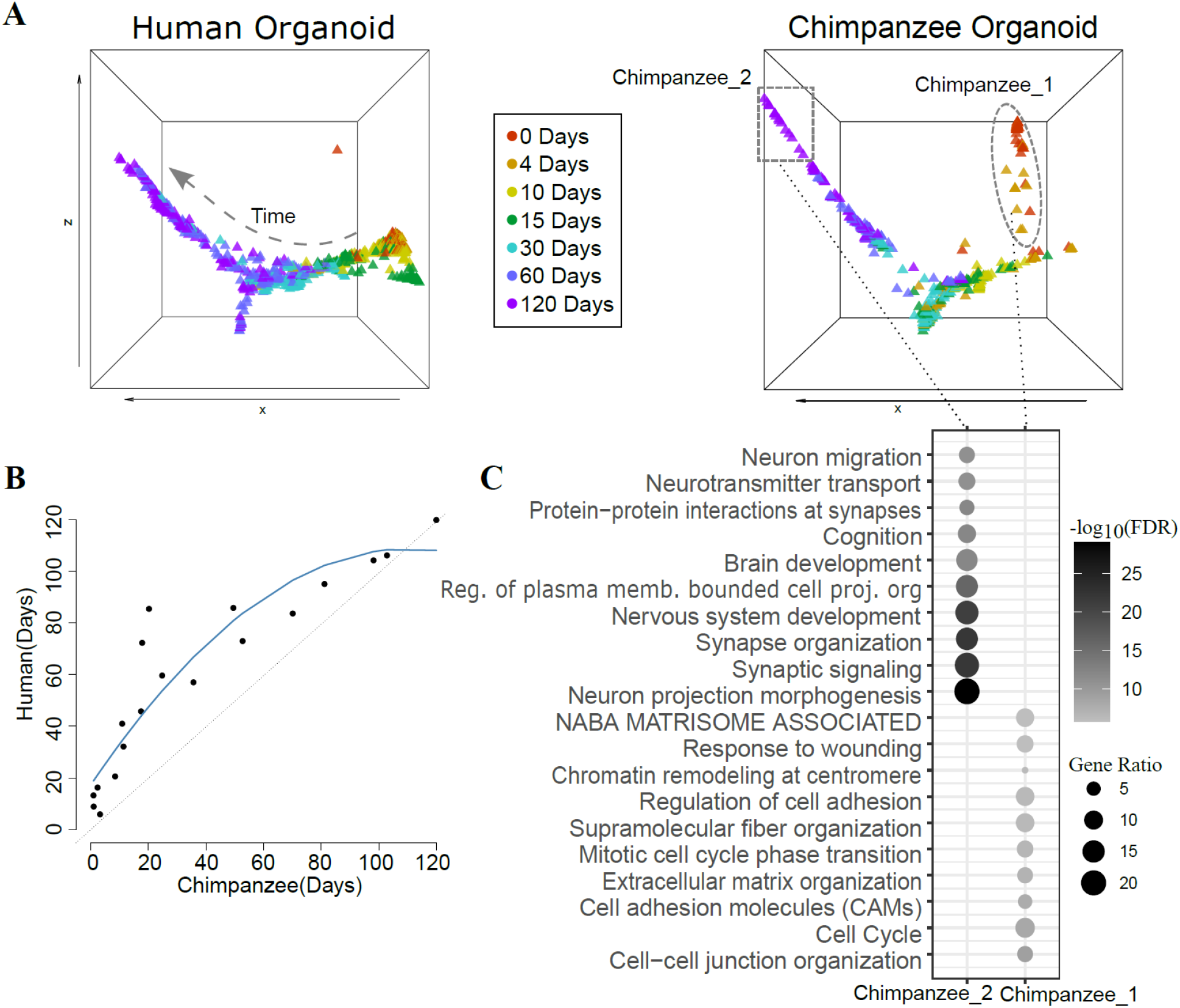
Developmental gene expression alignment between human and chimpanzee organoids. (**A**) The samples of human and chimpanzee organoid cells^33^ (visualized by pseudo-cells) in the common space after BOMA alignment. Human and chimpanzee organoid cells were plotted separately for comparison. The dot colors represent the experimental time points. The dashed line on the left panel shows the direction of the developing trajectory. Two chimpanzee organoid-specific clusters (Chimpanzee_1 and Chimpanzee_2) are highlighted on the right panel. (**B**) Timepoints matching between human and chimpanzee organoids. The aligned trajectories were cut into 30 segments along the x-axis. The sampling time of cells within each segment was averaged for human and chimpanzee organoids separately. Segments including both species show a matching timepoint in the plot, while segments with only one species were ignored. (**C**) Functional enrichments of the chimpanzee-organoid specific clusters genes.

Therefore, our BOMA alignment revealed a developmental gene expression similarity between human and chimpanzee organoids. Our analysis also uncovered the cross-species discrepancy in neural development and cellular functions. These results demonstrate the capability of BOMA to compare emerging organoid single-cell data and provide insights into underlying cellular and molecular mechanisms driving neurodevelopment.

### Cell-type level conservation in development between human brains and organoids derived from embryonic stem cells

To broaden BOMA applications to single-cell data sets of human brains versus organoids, we first benchmarked BOMA on two particular single-cell data sets. The comparison of the human brains and organoids, especially at the cell-type level, will greatly advance our understanding of how well *in-vitro* cultured organoids model the *in-vivo* human brain. We first aligned single-cell data from postmortem human brains (N=4,261, Dataset 3)^39^ with those of organoids differentiated from a well-established human embryonic stem cell (ESC) line (H9, N=11,048, Dataset 7, Table S1)^33^. This human brain dataset covers prenatal samples across 6-32 PCWs, while the organoid dataset includes organoids from 0 day up to 4 months *in vitro*. Before performing the alignment, human brain pseudo-cells (N=490) and organoid pseudo-cells (N=497) were generated from both datasets (Methods).

Based on our earlier analysis of bulk RNA-seq datasets (Figure 2B), organoids up to 4 months could be aligned across prenatal developmental periods, indicating that the developmental time ranges of these two separate studies are comparable. Thus, we aligned these two scRNA-seq datasets and identified five cell clusters in the common space (Methods). Each cluster represents a group of cells that likely have similar functions (Figure 4A). Interestingly, each individual cluster contains the cells from both human brains and organoids, suggesting that the *in-vitro* organoids likely is composed of the major cell types in the human brain. To understand functions underlying those clusters, we calculated the cell-types enriched in each cluster. For a given cluster, the enrichments were performed for the cluster cells from human brains and organoids separately. We observed that the enriched cell types in each cluster were generally matched between human brains and organoids (Figure 4B). For instance, Cluster 1 was mainly enriched for early developing cells, such as radial glia (RG), and oligodendrocyte progenitor cell (OPC); Cluster 2 was associated with intermediate progenitor cells (IPC), and newborn excitatory neuron (nEN); Cluster 3 was mainly mapped to excitatory neuron (EN); Cluster 4 was mainly mapped to inhibitory neuron (IN); Cluster 5 was mainly mapped to endothelial cells. The details of cell-type annotations can be found in Table S3.

**Figure 4.**
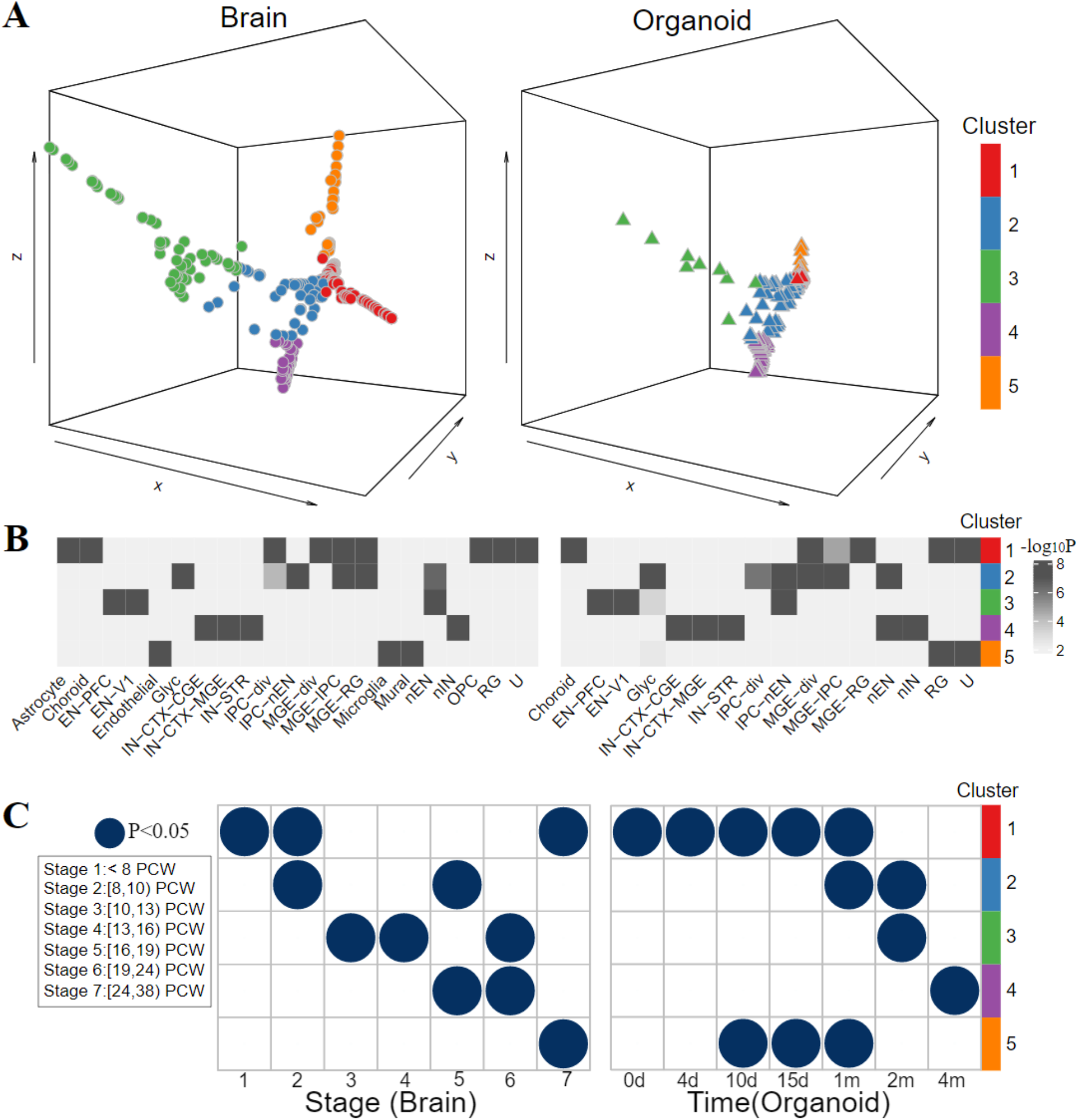
Alignment of developmental gene expression between human brains and human ESC organoids. (**A**) The human brain^39^ and organoid^33^ cells (visualized by pseudo-cells) on the common space after BOMA alignment. Left: human brain. Right: human organoids. Aligned cells were grouped into 5 clusters (by colors, Methods). (**B**) The associated cell-types of each cluster from the enrichment analysis (by hypergeometric test, Methods). ComplexHeatmap^40^ was used to plot the significance of associated cell-types for each cluster. (**C**) The associated developmental stages (time points) of cell clusters (by hypergeometric test, Methods). PCW: Post Conception Weeks. Dots represent associations with Benjamini-Hochberg (BH) adjusted p-values <0.05.

Moreover, it is also important to look at matching developmental timing between human brains and organoids. Determining the corresponding developmental periods during which cells are generated and specified in organoids will greatly benefit the design of culturing experiments. To address this, we identified the associated developmental stages of each cluster by calculating the significance of cells overlapping between each stage versus each cluster (Methods). In general, we observed each cluster was associated with different developmental stages (Figure 4C). Together with the fact that clusters are composed by different cell types, this indicated the dynamic maturation of cell types across development. Interestingly, we observed that brains and organoids development follow a similar pattern, which again supports the developmental conservation between two datasets.

However, discrepancies were observed for Cluster 5, which is significantly associated with microglia, endothelial and mural cells in the human brains, but with radial glia (RG, Table S3) in the organoids (Figure 4B), reflecting the fact that these brain cell types have distinct origins from neurons and glia in the brain. Also, Cluster 5 is associated with later developmental stages in brains (>24 PCWs) but with earlier cultured time-points in organoids (<1 month, Figure 4C). This suggests that human brain cells in this cluster are more mature than organoid cells. This observation was supported by BOMA-aligned cells on the common space in Figure 4A, where human brain cells stretched longer in this cluster than organoid cells.

### Large-scale alignment of integrated datasets in human brains and organoids derived from induced pluripotent stem cells

Brain organoids differentiated from iPSC have been used extensively to model human brain development and developmental disorders^18,41,42^. Here, we tested BOMA performance for aligning large-scale datasets of human brains versus both iPSC and ESC derived organoids. In particular, we integrated scRNA-seq datasets from multiple studies to align single cells of human brains and human brain organoids (Methods). The integrated datasets have 57 human brain samples and 28 iPSC or ESC derived organoids. The human brain data contains 175,334 cells across 5.85-37 PCWs, while the organoids data contains 187,179 cells across 21-105 cultured days. Similar to previous analyses, we first clustered cells into pseudo-cells (1,018 in human brains, 872 in organoids) to remove stochastic noise, and afterward evaluated the batch effects across datasets. The tSNE plots show that minimum batch effects persist after our reducing cells to pseudo-cells (Figure S3A,B; Figure S4A,B). We then input pseudo-cells into BOMA for alignment. We found that BOMA aligns the two large-scale integrated datasets reasonably well, showing aligned cell trajectories with similar cell-type distributions between brain and organoid cells in the common space (Figure 5A, Figure S3D, Figure S4D). For instance, OPCs were embedded in the middle, excitatory neurons, inhibitory neurons, and radial glia were aligned in a separate branch, while IPCs spread across both excitatory neurons and radial glial cells branches. Expectedly, even less batch effects were observed after BOMA alignment (Figure S3A,C, Figure S4A,C).

**Figure 5.**
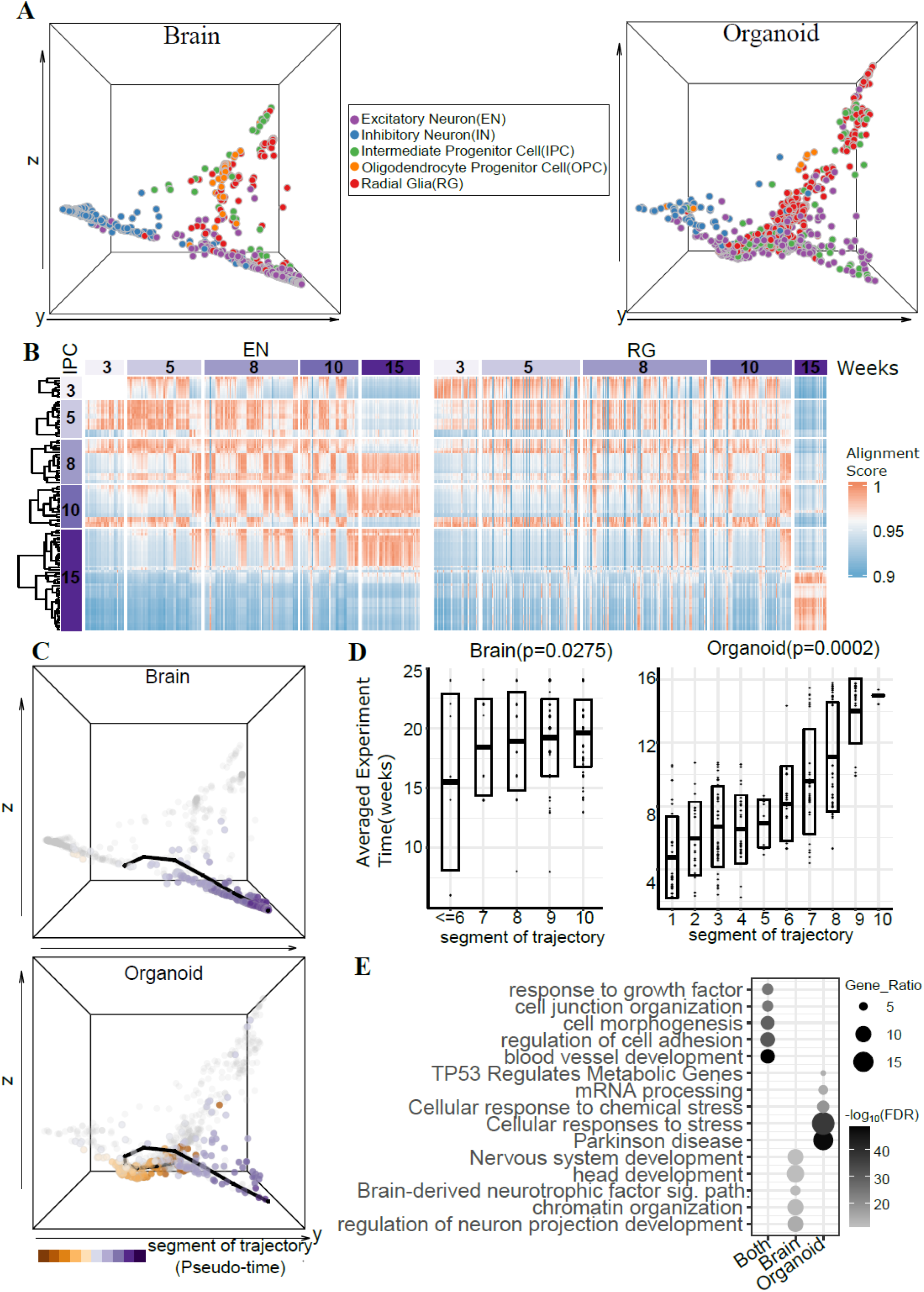
Large-scale alignment of integrated scRNA-seq gene expression datasets in human brains and organoids from multiple studies. Five scRNA-seq datasets of human brains (Nowakowski, T.J. et al. Science, 2017^39^; Trevino A.E. et al., Cell, 2021^45^; Bhaduri, A., et al., Nature, 2020^18^) and organoids (Birey et al., Nature, 2017^42^; Bhaduri, A., et al., Nature, 2020^18^) were applied (Methods). (**A**) The human brain and organoid cells (visualized by pseudo-cells) on the common space after BOMA alignment. Left: human brains. Right: organoids. The dots are colored by given cell-types from the datasets (Methods). (**B**) Experimental time correspondence between aligned intermediate progenitor cells (IPC) versus excitatory neurons (ENs)/radial glia (RGs) within organoids samples. (**C**) Inferred developmental trajectory for ENs based on their coordinates on the common space. Top: human brains. Bottom: organoids. (**D**) Trajectory segments vs. prior development stages (experimental timepoints). Human brain cells from segments earlier than Stage 6 were grouped together due to the limited sample sizes. Mann-Kendall trend test for mean values of each segment was used to test the trending significance (Methods). (**E**). The enriched functions and pathways of genes significantly upregulated in organoid ENs, genes upregulated in brain ENs and genes expressed in both brains and organoids.

Progenitor cells, such as IPCs, can divide and differentiate into postmitotic excitatory neurons in the developing cerebral cortex. This suggests that IPCs should align with neurons on the same maturational trajectory. To test whether this is true, we compared the developmental distribution of cultured IPCs with aligned excitatory neurons and radial glia within organoid samples. Interestingly, we did observe a time shift between IPCs with excitatory neurons (e.g., IPC of 3 weeks can align with EN of 10 weeks, IPC of 8 or 10 weeks can align with EN of 15 weeks, etc.), but not between IPCs and radial glia (Figure 5B).

Moreover, we benchmarked other state-of-the-art methods and compared them with BOMA. Although Seurat^16^ (Figure S5) and Liger^43^ (Figure S6) can perform alignment at single-cell level, both failed to identify the developmental trajectories. MetaNeighbor^44^, a correlation-based method for characterizing cell-type replicability across scRNA-seq datasets, had computational memory issues when applied on all cells within this dataset, and was unable to identify cell-type replicability on a 10% sub-sampled dataset (Figure S7). Several other manifold based alignment method (UnionCom^27^, SCOT^28^, MMD-MA^26^) can map the pseudo-cells into a manifold space, but the cell types were not embedded closely (Figures S8-10). In summary, BOMA outperforms other platforms in terms of both finding aligned cell trajectories and discovering cell-type developmental conservation across large-scale human brain and organoid datasets.

### Brain-organoid aligned trajectory analysis reveals conserved and distinct developmentally expressed genes in specific cell types

Aligned cell trajectories by BOMA between human brains and organoids show developmental processes across various cell types. To further understand the gene expression programs driving cell-type maturation, we identified maturation trajectories based on the coordinates of cells corresponding to each cell type in the common space, such as excitatory neurons (Figure 5C) and IPCs (Figure S11). Then, we identified the genes that are differentially expressed (DEGs) across the cell-type trajectory between human brains and organoids. The enrichment analysis of those DEGs revealed conserved and specific developmental functions of the cell type across human brains and organoids (Methods).

The cell-type trajectories revealed the pseudotimes of individual cells during development (i.e., cell positions over the trajectory), hypothetically providing higher timing resolution than the prior timing information (Methods). By cutting the trajectory into segments and correlating them with the developing stages, we found that the segments of such pseudo-times significantly correlate with real developmental stages (Figure 5D for EN with adjusted p<0.0275 in brains, p<0.0002 in organoids, and Figure S12 for IPC trajectory), which suggests that this trajectory (pseudo-time) captures the real developmental maturation of cell types.

We then identified the DEGs for each segment along each cell-type’s trajectory (Methods). We identified 549 organoid and 310 brain upregulated genes that were differentially expressed within at least one segment of the excitatory neuron’s trajectory (Data 2). Functional enrichment of these DEGs showed that organoids upregulated genes were mapped to chemical stress response, which is supported by a previous study^18^(Figure 5E). On the other hand, the brain upregulated genes were mapped to brain development processes, as expected. To validate the differential expression of some of these genes, we performed immunofluorescence in the developing human neocortex and human organoids at different stages of differentiation (Figure S12). We found that the expression changes of important genes across stages (percentage of expressed cells) are greatly consistent with our results.

*SATB2*, encoding a transcription factor shown to modulate cortical neuron projection identity^46^ displayed no expression throughout the maturation of human organoids, whereas *PSMB5*, encoding a 20S proteasome subunit, demonstrated consistent expression across the maturation trajectories by BOMA (Figure 6A). These findings were confirmed with immunofluorescent staining of human organoids of *SATB2* (Figure 6C, <1.5% cells) and *PSMB5*(Figure 6I, ∼10% of cells) across 8-weeks, 10.5-weeks, and 14-weeks differentiation time points corresponding to segments 7, 8, and 9, respectively (Figure 6C and 6I). Also, *POU3F2*, encoding a transcription factor important for primate radial glia expansion and differentiation^47^, displayed no expression throughout organoids maturation by BOMA (Figure 6A), and indeed it was found expressing in a small number of cells (∼3%) in organoids by immunostaining (Figure 6F).

**Figure 6.**
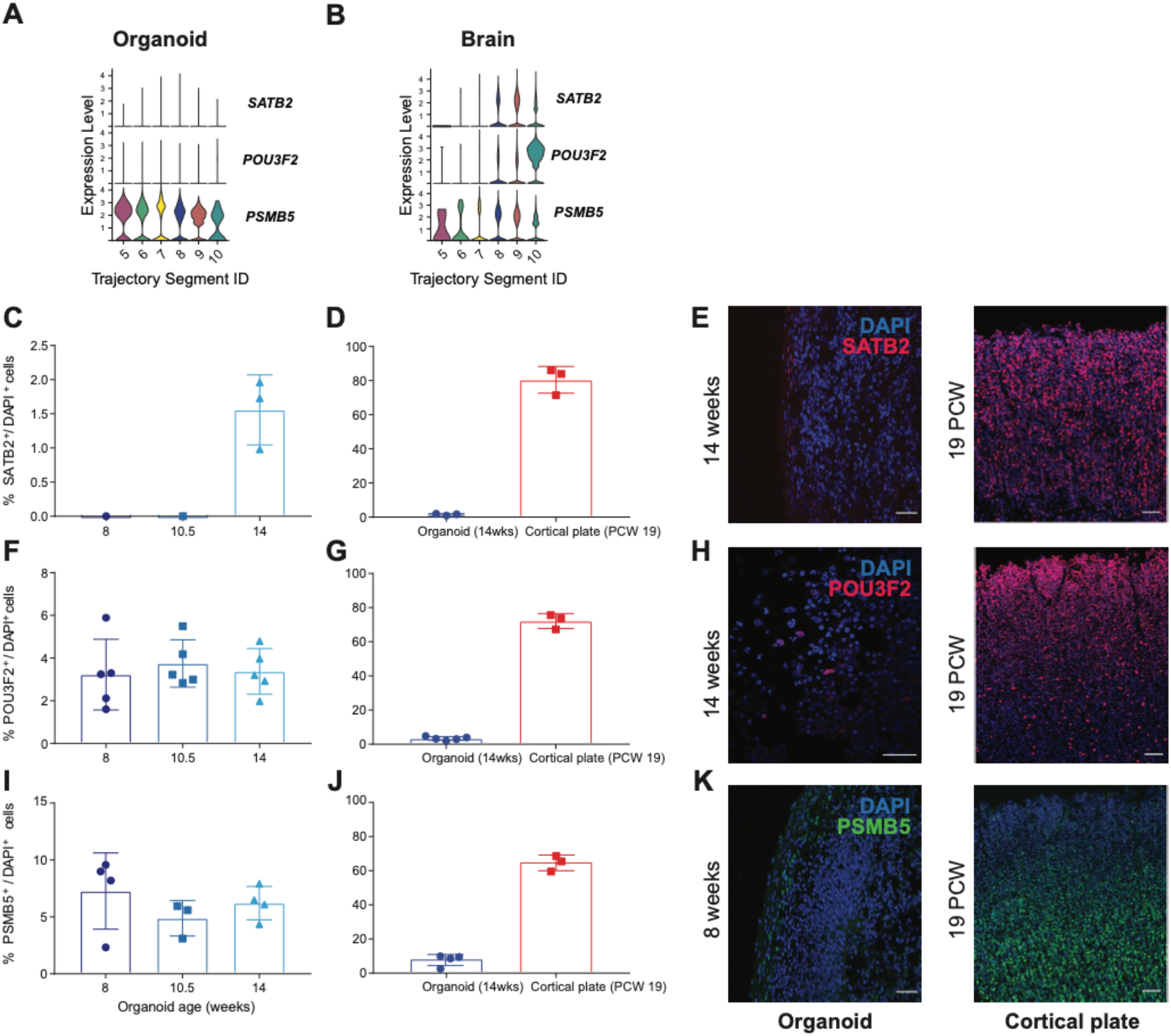
Experimental validation of developmental expression of predicted brain- and organoid-specific genes. (**A, B**): Developmental expression profiles of *SATB2, POU3F2*, and *PSMB5* mRNAs in human organoids (**A**) and human neocortex (**B**), determined by BOMA. Time correspondence for each segment ID can be found in **Figure 5D**. (**C, F, I**): Immunostaining of cortical organoids (n=3) revealed percentages of cells expressing *SATB2*(**C**), *POU3F2*(**F**) and *PSMB5*(**I**) during the maturation at 8, 10.5 and 14 weeks. (**D, G, J**): Quantification of SATB2^+^, POU3F2^+^, and PSMB5^+^ cells showed significant enrichment for human cortical plate (PCW 19, correspond to segment 9) as compared to organoids (14 weeks, corresponding to segment 9). Differences between organoid and cortical plate were tested using unpaired t-test with Welch’s correction. *p*<0.0038 for SATB2^+^ cells, *p*<0.001 for POU3F2^+^ cells, and *p*<0.0011 for PSMB5^+^ cells. (**E, H, K**): Representative images of organoid and human brain sections. Scale bar: 50 μm.

On the other hand, *SATB2* and *POU3F2* were identified as significantly upregulated in excitatory cells of the human neocortex as compared to human organoids at late stages (∼19PCW) by BOMA (Figures 6A-B, BH adjusted Wilcoxon rank sum test *p*<1e-2). Immunofluorescent staining confirmed the enrichment of SATB2^+^ and POU3F2^+^ cells in the human cortical plate of 19PCWs as compared to human organoids of 14 weeks (Figures 6D-E with t-test *p*<0.0038 and 6G-H with t-test *p*<0.001). In addition, *PSMB5* expressed higher than two other genes at earlier stage in both organoids and brains but comparatively only in brains at later stage by BOMA (Figures 6A-B, which is also observed in Figures 6D, 6G, 6J. Significant more PSMB5^+^ cells were found to be enriched in human neocortex (t-test *p*<0.0011), but mostly in deep layer and subplate neurons (Figure 6K), suggesting that human organoids may have lower abundance of subplate neurons.

## Discussion

In this work, we present BOMA as a framework for comparative analysis of gene expression between brains and organoids, with an attempt to understand the genomic regulations during their development. Our evaluation of BOMA on both bulk tissue and single-cell datasets demonstrated its scalability. Spatiotemporal and species-wise gene expression patterns have been observed by our alignment. Genes differentially expressed across cell-types and developmental stages were also identified by our scRNA-seq analysis. Although we only focused on comparing RNA-seq datasets between brains and organoids, BOMA can be easily applied to compare pairs of any samples (RNA-seq or other modalities). Hence, we provide a web tool of BOMA for general community use.

scRNA-seq data are in general noisy and stochastic, the pseudo-bulk method we benchmarked demonstrated these approaches can diminish scRNA-seq noises as well as balance sample sizes across datasets. The future development of more accurate scRNA-seq technologies will potentially improve the alignment of single cells. Also, the scRNA-seq datasets we analyzed were integrated from multiple published studies, so the input of BOMA can be confounded by various experimental factors, for instance, the sample-wise batch effects, the organoids culturing periods, the sample sizes, sample time, and sequencing depths, etc. As we showed in the results, BOMA significantly reduced these experimental confounders and demonstrated superior performance for integrative analysis of multiple studies. However, our evaluation was only based on limited samples from limited cultured periods, with limited numbers of pseudo-cells. Future studies using longer cultured organoids and more samples are recommended for better comparative analysis between brains and organoids. In addition, as a framework, BOMA can easily incorporate other existing alignment methods (e.g., Manifold Warping, CCA, etc.). For designing the framework, we also pursued a supervised manner, which allows the correspondences between sample (cell) pairs to be incorporated into the alignment as prior knowledge. For example, users can define any correspondence information based on their own domain expertise; Also, cell correspondences generated by other alignment tools (e.g., Seurat integration, Liger, etc.) can also be incorporated as prior knowledge of BOMA by defining the correspondence matrices.

Our manifold alignment analysis showed gene expression similarities between organoids and brains, demonstrating the viability of using organoids to understand human brain development^15^. However, differences were also observed in the comparative analysis, which suggests future protocol optimizations are needed^9^. Previous studies have demonstrated the wide application of organoids as experimental models for drug-screening of neuro-diseases^13^ and other diseases, e.g., cancer^48,49^. Other studies have also shown using patient-derived organoid (PDO) platforms to improve preclinical drug discovery in personalized medicine^50^. Recent clinical trials are moving towards cell therapy of diseases using lab cultured organoids^51–53^. All these reports suggested unprecedented opportunities for organoids in both lab research and clinical treatment. And we believe our comparative analysis framework, BOMA, which allows a deeper understanding of the gene regulatory mechanisms underlying the cultured organoids, will benefit future clinical studies.

## Methods

### Brain-Organoid Manifold Alignment (BOMA)

Emerging organoids have been widely used as models to mimic complex brain development. We developed BOMA pipeline to use manifolds to align gene expression data between brain and organoid samples (e.g., tissues, cells) (Figure 1). Such brain-organoid expression data alignment from BOMA aims to uncover conserved (aligned) and specific (unaligned) developmental gene expression patterns across brains and organoids. Our further downstream analyses of such expression patterns allow a deeper understanding of developmental functional genomics at both tissue and cell-type levels, especially in organoids.

Suppose that we want to compare two developmental gene expression datasets (e.g., brains vs. organoids) matrices, *X* = [*x*_1_, *x*_2_, …, *x*_*m*_] ∈ *R*^*d*×*m*^ and *Y* = [*y*_1_, *y*, …, *y*_*n*_] ∈ *R*^*d*×*n*^, where *d* is the number of genes, *m* and *n* are the number of samples within each dataset, *x*_*i*_ ∈ *R*^*d*^ is a *d* - dimensional vector representing the expression levels of *d* genes in the *i*^*th*^ sample in *X*, and *y*_*j*_ ∈ *R*^*d*^ is also a *d*-dimensional vector representing the expression levels of *d* genes in the *j*^*th*^ sample in *Y*. The samples of {*x*_*i*_, *i* = 1,2, …, *m*} and {*y*_*j*_, *j* = 1,2, …, *n*} are ordered by prior timing information if available. BOMA carries out the alignment by two major steps. In Step 1, BOMA globally aligns brain and organoid samples, based on prior timing (or any sequential) information of samples. Such prior timing information is typically at low resolution, e.g., only cultured days available for many cells in organoids. This global alignment establishes the initial correspondence across brain and organoid samples. In Step 2, from such initial correspondence, BOMA applies manifold learning to locally refine the alignment and co-embed brain and organoid samples onto a common manifold space. The manifold shapes of the samples on the space are expected to uncover various developmental trajectories, which can be either conserved across brains and organoids (aligned samples) or brain/organoid specific (unaligned samples). Furthermore, the manifold shapes from the space are expected to form developmental trajectories, revealing potential pseudo times among samples. Such pseudo times, at a refined high resolution, provide unobserved timing from prior information.

#### BOMA Step 1 - Global Alignment

This step aligns X and Y at a coarse-grained level and initializes the correspondence matrix (*W*) for the next step. Primarily, we introduce two popular methods for global alignment.

1. Dynamic Time Warping (DTW). DTW finds the optimal set(s) of aligned samples (*π*\*) between *X* and *Y* by minimizing the sum of distances between all aligned sample pairs:

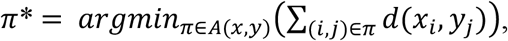

where *d*(*x*_*i*_, *y*_*j*_) is the distance between the i^th^ and j^th^ samples of X and Y, *A*(*x,y*) is the set of all possible alignments between the two datasets. Distance of the samples x and y used here is defined by 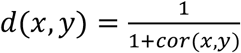, where cor(*x, y*) is the Pearson Correlation. Specifically, we used R package, dtw^54^ to perform the DTW alignment. In this work, we chose the constraint as ‘open begin and end’, which means that two sequential datasets can be unaligned at the beginning and end. The aligned samples from DTW can be used to initialize a corresponding matrix among samples, *W*, where *W*_*ij*_=1 if samples *x*_*i*_, *y*_*j*_ are aligned, and = 0 otherwise.
2. Correlation based kNNgraph: This method first calculates the Pearson Correlation of each sample pair and then constructs a k-nearest neighbor graph (kNNgraph) by linking each sample with its k (a hyperparameter) most correlated neighbors in the other dataset. The adjacency matrix of the constructed kNNgraph can thus be used as the correspondence matrix *W*.

Besides the two methods above, this step can also be accomplished by other methods, e.g., Liger^43^, which uses Nonlinear Matrix Factorization(NMF) for single-cell alignment; Seurat^16^, which aligns single cells by identifying anchor genes.

#### BOMA Step 2 - Local Alignment

this step performs a manifold alignment of *X* and *Y* using the correspondence matrix (*W*) from Step 1 as the initial alignment. Specifically, manifold alignment finds shared manifolds of samples from *X* and *Y* and maps them onto a common space. The proximate samples on this space suggest well aligned, whereas distant samples for unaligned. To this end, it aims to find the functions *f*_*X*_^*^ and *f*_*Y*_^*^ that minimize the following loss function to map the samples onto the common space:

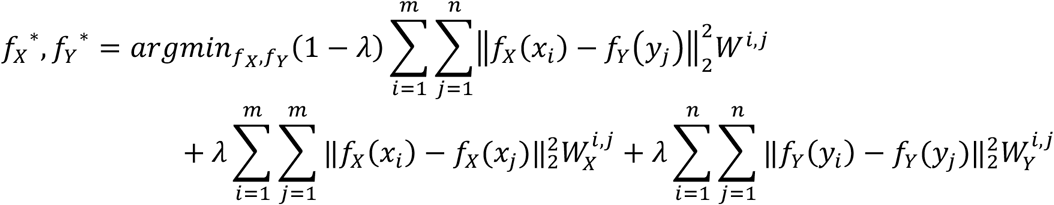

where *W*^*i,j*^ is the correspondence between *x*_*i*_ and *y*_*j*_ from Step 1. It can be weighted, or it can be binary (e.g. 0: aligned, 1: un-aligned) as in this work. *W*_*X*_ and *W*_*Y*_ are two neighborhood similarity matrices, which were generated by kNNgraph. *λ* is a scalar, which constitutes the trade-off between the alignment across datasets and preserving manifolds within datasets. By default, we set *λ* equals 0.5. Here, we use nonlinear manifold alignment (NMA) to solve the above optimization problem. NMA is non-parametric and directly estimates the coordinates of samples on the common space from optimal alignment via eigen-decomposition^24^. Also, we implement NMA in this step using our previous method and tool, ManiNetCluster^24^.

After mapping samples onto the common manifold space by BOMA, we can calculate the Euclidean distances of samples on the space, i.e., *d*_*ij*_ for Samples *i* and *j*. An alignment score between samples *i* and *j* can be then defined by 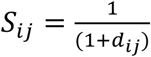. The high alignment scores suggest the well aligned samples.

### Gene expression datasets of brains and organoids

As summarized in Table S1, we collected recently published RNA-seq gene expression datasets for brains and organoids, covering both bulk tissues and single cells across differential developmental stages.

Briefly, <u>Dataset 1</u>^4^ contains bulk-tissue RNA-seq of 826 samples from 16 regions of human brains (n=460) and 9 regions of RM brains(n=366) in Brainspan and PsychENCODE projects. <u>Dataset 2</u>^55^ contains single-cell RNA-seq (scRNA-seq) of 40,000 cells from human brain germinal zone and developing cortex regions between 17-18 Postconceptional Weeks (PCWs). <u>Dataset 3</u>^39^ contains scRNA-seq data of 4,261 cells in human brains between 6-32 PCWs. <u>Dataset 4</u>^45^ contains scRNA-seq data of 57,868 cells from four human brain primary samples at different developmental stages between 16-24 PCWs. <u>Dataset 5</u>^18^ includes scRNA-seq of 136,254 cells from human brain samples collected at 14,18,22 PCWs. <u>Dataset 6</u>^11^ is from the cultured organoid samples and includes bulk RNA-seq data of 62 samples from ten time points between 50 days to ∼ two years. <u>Dataset 7</u>^33^ contains scRNA-seq of 73,358 cells in organoids from human or chimpanzee between 0 days to four months. <u>Dataset 8</u>^42^ contains scRNA-seq of 11,838 cells from organoids cultured for 105 days. <u>Dataset 9</u>^18^ contains scRNA-seq of 189,346 organoid cells of culturing time spanning 3-10 weeks.

### Identification of human brain developmental genes

We identified a set of genes related to human brain development at both tissue- and cell-type levels, as input features for BOMA alignment (Figure S13). First, we used the bulk RNA-seq data in BrainSpan^4^ to predict co-expression gene modules within each brain tissue (region) by WGCNA^56^. We identified 1,191 co-expression gene modules in total. Genes from the same module are co-expressed at certain tissue across the development, suggesting that they are likely co-regulated and thus involved in similar biological processes, so we term them as ‘development modules’. Second, we applied Scanpy^57^ on the single-cell RNA-seq dataset from Dataset 2^58^ (Table S1) to identify developmental expressed genes at the cell-type level. Specifically, for each of 11 cell-types, we compared this cell-type with all other cell-types and found cell-type differentially expressed genes (adjust p-value < 0.01 and log fold change > 1). After iterating through 11 cell-types, we identified 2,032 cell-type expressed genes in total. Third, we overlapped each development module from a tissue-type with each cell-type expressed gene set and performed hypergeometric tests (R function phyper()) to determine the significance of the developmental gene overlaps between the tissue-type and the cell-type (tissue-cell-type pair). We also adjusted the p-values of tests using ‘Benjamini-Hochberg procedure (BH)’. We selected the overlapped genes of tissue-cell-type pairs with adjusted p<0.01 as significant overlapped gene sets. Finally, we obtained 1,533 genes as human brain developmental genes (Data 3).

### scRNA-seq data pre-processing

We used Seurat^16^ to preprocess all applied scRNA-seq datasets. In particular, we removed the cells expressing less than 200 genes and the genes expressed within less than 30 cells. The rest cells were filtered by mitochondrial genes to be less than 10. The preprocessed datasets were then log2 transformed.

Compared to bulk RNA-seq, scRNA-seq is noisy with random effects. To address this, recent studies^59,60^ used pseudo-bulk methods to aggregate single-cells across biological replicates and improved downstream differential expression gene analyses. Therefore, we also applied the pseudo-bulk methods^33,59,60^ to create pseudo-cells from single cells. Specifically, we first grouped single cells into cell clusters. Each cluster represents one pseudo cell, and its expression levels are the averaged gene expression of cells within the cluster. This step can also balance the sample sizes across datasets, e.g., numbers of pseudo cells.

In particular, we benchmarked two major pseudo-bulk methods, PCA-based^33^ and Seurat, and found the one for each application leading to a better BOMA alignment. The PCA based method calculated the principal components (PCs) of single cells, and then hierarchically clustered (R function ‘hclust’) single-cells with the top 20 PCs to generate pseudo-cells, i.e., cell clusters. We used the function FindClusters() in Seurat for clustering single-cells as Seurat-based method. We used the PCA-based method for the analyses in Figure 3 and Figure 4, which were consistent with the paper generating the data^33^. However, we benchmarked the latter method and found it works better than the PCA-based method, so we used the Seurat-based method for the analysis in Figure 5.

To determine how the alignment is affected by the number of pseudo-cells, using the dataset of Figure 5, we tested different numbers of pseudo-cells by adjusting the ‘resolution’ parameter in the Seurat FindClusters() function. In this way, we generated pseudo-cells that varied from ∼1,000 to ∼10,000 (Figure S14A). To evaluate the alignment accuracy, within the aligned common manifold space, we calculated the pairwise distances of pseudo-cells of the same cell-type. Specifically, the coordinates of pseudo-cells were standardized per pseudo-cell, then distances between pseudo-cells of the same cell-type were averaged. Interestingly, the experiment result shows BOMA is scalable to the number of pseudo-cells, with the pairwise distances not significantly affected (Figure S6B). Considering this characteristic, and in order to balance the number of pseudo-cells across datasets, we set a lower resolution for datasets with more cells and set a higher resolution for datasets with fewer cells for the later analysis (Table S1). In particular, for organoid data, we set the resolution values as 10 for Dataset 8 and 1 for Dataset 9; for brain data, we set the resolution values as 10 for Dataset 3, 5 for Dataset 4 and 1 for Dataset 5.

### Gene set enrichment analysis

We used Metascape^61^ to perform the gene set enrichment analysis. The enriched categories include KEGG pathways, Gene Ontology (GO) terms, protein-protein interactions, and diseases (via DisGeNET). The false discovery rates (FDRs, q-values) were used to quantify the enrichment significance.

### Clustering BOMA-aligned samples and differentially expressed genes of clusters

We applied the Spectral clustering from Python package ‘sklearn’^62^ to cluster aligned samples on the common space, based on their alignment scores. We also identified differentially expressed genes (DEGs) of clusters. To this end, we used Presto^63,64^ to perform the Wilcoxon rank sum test and auROC analysis by comparing cells from each cluster with all others cells in the dataset.

### Harmonization of cell types across datasets

Cell type names may vary across studies. For instance, cell types from Organoid Dataset 7 are broad and different from many refined types in human brain. To solve this, we reassigned the types of the cells in Dataset 7 using the human brain cell-types in Dataset 3 by the ‘TransferData’ function in R package Seurat^16^. We also merged some sub-cell-types to their broader types, e.g., EN-PFC1 to EN-PFC. Besides, even different studies for brains or organoids can have different sub-cell-types. To make cell types across these studies comparable, we grouped annotated cell-types from each study into common major cell-types (Table S4) for downstream comparative analyses. Also, for Dataset 8 without cell-type information, we annotated cell types using known cell-type marker genes^42^ with Seurat.

### Hypergeometric enrichment of cell-types and developmental time stages

For the cell clusters from BOMA applications to single cell data, we calculated their cell-type enrichments (or developmental timepoint enrichments), revealing possible cellular and developmental functions of the clusters. In particular, a hypergeometric test was performed for such enrichment analysis, with the p-values being calculated as:

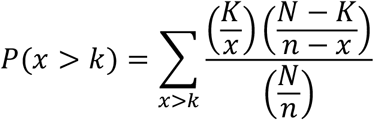

where *N* is the total number of cells, *n* is the total number of cells of a certain cell-type (or cells from a certain developmental timepoint), *K* is the number of cells in the cluster and *k* is the number of cluster cells of certain cell-type (time point) in the cluster. Finally, we corrected the p-values using BH method and selected p<0.05 as a significant threshold for enrichments.

### Trajectory analysis for BOMA alignment

Since BOMA applies the manifolds to align single cells between brains and organoids, the manifold shapes from aligned cells are expected to reveal potential developmental trajectories. To further identify such trajectories, we used SCORPIUS^65^ to infer the developmental trajectory for each cell-type on the common space. Primarily, for each cell-type, we input the 3D coordinates of its cells on the common space from BOMA to the infer_trajectory() function of SCORPIUS (maximum iteration of 100) to output the trajectory. To determine a root on the trajectory, we first cut the trajectory into 10 continuous segments. Each cell was assigned to the closest segment based on the distance. Then a developmental time for each segment can be determined by averaging the prior times of all cells in that segment. We then assign the segment with minimum averaged time as the root. Besides, for each cell type, we also used FindMarkers() in Seurat^16^ to identify differentially expressed genes in the type’s cells of each segment, i.e., “Segment cell-type DEGs” implying development-stage-specific gene expression patterns at the cell-type level. To allow gene expression values to be comparable across datasets, the IntegrateData() function of Seurat was used to integrate datasets by identify a set of anchor genes.

### Experimental validation of genes in specific cell types and developmental stages

Human WC5907 iPSC line^66^ was maintained on mouse embryonic fibroblast feeder layers as described^67^ and differentiated into organoids as carried using a published protocol^68^. Briefly, iPSCs were lifted using dispase (0.4mg ml^-1^) and transferred to low attachment flasks (Greiner Bio-One) in hESC media plus two SMAD inhibitors SB-431542 and LDN-193189 from days 0-5. Organoids were then switched to neural medium plus growth factors EGF (20ng ml^-1^; R&D Systems) and FGF2 (20ng ml^-1^; WiCell) from days 6-24. After 24 days, organoids were cultured in neural medium supplemented with growth factors BDNF (20ng ml^-1^; Peprotech) and GDNF (20ng ml^-1^; Peprotech) until day 43 with media changes every 2-3days. Organoids were collected at 8, 10.5, and 14 weeks of differentiation and fixed with 4% PFA overnight. They were then washed with PBS 3x for 15min, and transferred to a 30% sucrose solution for 48hrs. Organoids were embedded in OCT and 30% sucrose (1:1) and stored in -80 freezer until analysis.

Fixed organoids were cryosectioned (17um) and stained with antibodies against proteins and markers of interest as described^68^. Organoid sections were washed with PBST (PBS containing 0.1% Triton X-100) and blocked in blocking buffer (10% normal goat serum (Sigma-Aldrich) and 0.3% Triton X-100 in PBS) for 1 hour at room temperature. Primary antibodies - anti-BRN2 (mouse, 1:500, Santa Cruz, SC-393324), anti-PSMB5 (rabbit, 1:1000, Novus Bio, NBP-13820), or anti-SATB2 (mouse, 1:100, Gen Way, 20-372-60065), anti-SOX2 (Mouse, 1:500, Abgent, Am2048a), anti-TBR1 (Rabbit, 1:1000, Abcam, Ab31940), or anti-CTIP2 (Rat, 1:500, Abcam, ab18465) were diluted in blocking buffer and incubated with the organoid sections overnight at 4°C. Sections were then washed 4 × 5 min with PBST. Alexa Fluor secondary antibodies (Thermo Fisher Scientific) were diluted in blocking buffer and incubated with organoid sections for 35min. at room temperature. Organoid sections were washed 4 × 5 min with PBST and counterstained with DAPI. They were then washed 2 × 5 min with PBST. Sections were scanned and visualized using either a Nikon A1 confocal microscope (Nikon) or an AxioImager Z2 ApTome microscope (Zeiss). The numbers of marker positive cells were quantified by unbiased stereology using StereoInvestigator software (MicroBrightField, Inc) as described^69^ PCW 19 human neocortex was fixed in 10% neutral buffered formalin at 4°C for 72 hours, cryoprotected with incubation in successive solutions of 10%, 20%, and 30% sucrose, and stored in 30% sucrose + 0.1% sodium azide. For validation experiments, PCW 19 human neocortex was embedded in Optimal Cutting Temperature (OCT) compound, cryosectioned at 30um thickness, and mounted on TOMO® adhesion slides (Matsunami Glass USA #TOM-11/90). Sections were washed in PBS (2 × 15 min) and incubated in blocking solution containing 5% (v/v) normal donkey serum (Jackson ImmunoResearch Laboratories) and 0.3% (v/v) Triton X-100 in PBS for 30 min at room temperature. Primary antibodies - anti-BRN2 (mouse, 1:500, Santa Cruz, SC-393324), anti- PSMB5 (rabbit, 1:1000, Novus Bio, NBP-13820) or anti-SATB2 (mouse,1:100, Gen Way, 20-372-60065) were diluted in blocking solution and incubated with tissue sections for 24 h at 4°C. Sections were washed with PBST (1X PBS + 0.3% Triton X-100) prior to being incubated with the appropriate fluorophore-conjugated secondary antibodies (Jackson ImmunoResearch Labs) for 30 min at room temperature. All secondary antibodies were raised in donkey and diluted at 1:250 in blocking solution. Sections were washed with PBST (3 × 5 min), treated with Autofluorescence Eliminator Reagent (Millipore #2160) according to manufacturer instructions, and coverslipped with Vectashield Plus Antifade Mounting Medium (Vector Laboratories #H-1000). Human neocortical samples were imaged on a Nikon A1 confocal microscope. Z-stack images taken at 20x magnification with a step size of 2um were imaged from n=3 sections. CellProfiler software was utilized to quantify positive cells. Difference significance between organoid and human cortical plate marker positive cell percentages was test by unpaired t-test with Welch’s correction.

## Author Contributions

D.W. designed and directed the study. C.H. performed research and analyzed data. A.M.M.S. and X.Z. designed the experiment validation. N.C.K. developed BOMA web app and provided customized scripts. S.O.S and R.R. performed experimental validation and interpreted experimental results. C.Y. benchmarked alignment tools. M.S. contributed to data analysis and the web app development. Q.C., A.M.M.S., X.Z. helped design the study. C.H., A.S., S.O.S, R.R., S.K., X.Z. and D.W. wrote and edited the manuscript. All authors read and approved the final manuscript.

## Acknowledgements

The authors would like to thank Biomedical Computing Group in Department of Biostatistics and Medical Informatics at University of Wisconsin at Madison for providing computing resources. We also thank the Intellectual and Developmental Disabilities Research Center in Waisman Center for valuable comments.

## Code availability

Codes, tutorial, and demo for BOMA are available at https://github.com/daifengwanglab/BOMA. A web app of BOMA is available at http://daifengwanglab.org/boma-webapp/.

## Data availability

All our results are provided in Supplementary Data (Data 1-3). All processed data are available at https://zenodo.org/record/6625362#.YqFxNufMKF4.

## Supplementary information

Data 1: marker genes of two chimpanzee organoids specific clusters

Data 2: differentially expressed genes within excitatory neurons of human brains or organoids

Data 3: 1,533 human brain development related genes

Supplementary Materials: Tables S1-4 and Figures S1-14

