## Supplemental Materials for "Brain and Organoid Manifold Alignment (BOMA), a machine learning framework for comparative gene expression analysis across brains and organoids"

### Supplementary

#### Supplementary Tables

##### Table S1. Summary of transcriptome datasets included in this study

| ID | Paper | Time Range* | Number of cells | Source | Use in this work |
|---|---|---|---|---|---|
| 1 | Li et al. 2018 | 8PCW – 40 Y | Bulk | brain | • Identify brain tissue-wise co-expression gene modules (development module)<br>• Bulk RNA-seq alignment (Figure 1) |
| 2 | Polioudaks, et al. 2019 | 17-18PCW | 40,000 | brain | Identify cell-type specific genes for brain development |
| 3 | Nowakowski, et al. 2017 | 6PCW – 32PCW | 4,261 | brain | Human brain single-cell dataset 1 for large dataset alignment (Figure 5) |
| 4 | Trevino et al. 2021 | 16PCW, 20PCW, 21PCW, 24PCW | 57,868 | brain | Human brain single-cell dataset 2 for large dataset alignment (Figure 5) |
| 5 | Bhaduri et al. 2020 | 14PCW, 18PCW, 22PCW | 136,254 | brain | Human brain single-cell dataset 3 for large dataset alignment (Figure 5) |
| 6 | Gordon et al. 2021 | 50D,75D,100D,150D,200D,250D,300D,350D,400D,600D | Bulk | organoid | Bulk RNA-seq alignment (Figure 1) |
| 7 | Kanton et al. 2019 | 0D, 4D, 10D, 15D, 1M, 2M,4M | 73,358 | organoid | • Cross-species single-cell alignment (Figure 3)<br>• Organoid single-cell alignment in Figure 4; |
| 8 | Birey et al. 2017 | 105D | 11,838 | organoid | Organoid single-cell dataset 1 for large dataset alignment (Figure 5) |
| 9 | Bhaduri et al. 2020 | 3W, 5W,8W,10W | 189,346 | organoid | Organoid single-cell dataset 2 for large dataset alignment (Figure 5) |

*Y: year; M: Month; W: week; D: day; PCW: Postconceptional Week

**Table S2. Number of Gene Modules identified by WGCNA for each brain region**

| Brain region | Number of Gene Modules |
|---|---|
| A1C | 33 |
| AMY | 92 |
| CBC | 114 |
| DFC | 96 |
| HIP | 53 |
| IPC | 68 |
| ITC | 37 |
| M1C | 17 |
| MD | 54 |
| MFC | 126 |
| OFC | 38 |
| S1C | 99 |
| STC | 95 |
| STR | 44 |
| V1C | 109 |
| VFC | 116 |

**Table S3. Interpretation of cell-types annotated in Figure 4B.**

| Cluster Name | Cluster Interpretation (Nowakowski, et al., 2017) |
|---|---|
| Astrocyte | Astocyte |
| Choroid | Choroid |
| Endothelial | Endothelial |
| EN-PFC | Early and Late Born Excitatory Neuron PFC |
| EN-V1 | Early and Late Born Excitatory Neuron V1 |
| Glyc | Glycolysis |
| IN-CTX-CGE | CGE/LGE-derived inhibitory neurons |
| IN-CTX-MGE | MGE-derived Ctx inhibitory neuron |
| IN-STR | Striatal neurons |
| IPC-div | Dividing Intermediate Progenitor Cells RG-like |
| IPC-Nen | Intermediate Progenitor Cells EN-like |
| MGE-div | dividing MGE Progenitors |
| MGE-IPC | MGE Progenitors |
| MGE-RG | MGE Radial Glia 1 |
| Microglia | Micrgolia |
| Mural | Mural/Pericyte |
| nEN | Newborn Excitatory Neuron |
| nIN | MGE newborn neurons |
| OPC | Oligodendrocyte progenitor cell |
| RG | Radial Glia |
| U | Unknown early cells |

**Table S4. Correspondence for sub-cell-types to be grouped into common major cell-types**

| Sub cell-type | Major cell-type |
|---|---|
| **Bhaduri et al. 2020 (organoid), Dataset 5** | |
| Astrocyte | Astro |
| ExcitatoryNeuron | EN |
| InhibitoryNeuron | IN |
| IPC | IPC |
| Outlier,Unknown | Unknown |
| RadialGlia | RG |
| **Nowakowski et al. 2017, Dataset 3** | |
| Astrocyte | Astro |
| Choroid | Choriod |
| EN-PFC1,EN-PFC2,EN-PFC3,EN-V1-1,EN-V1-2,EN-V1-3,nEN-early1,nEN-early2,nEN-late | EN |
| IN-CTX-CGE1,IN-CTX-CGE2,IN-CTX-MGE1,IN-CTX-MGE2,IN-STR,nIN1,nIN2,nIN3,nIN4,nIN5 | IN |
| IPC-div1,IPC-div2,IPC-nEN1,IPC-nEN2,IPC-nEN3,MGE-IPC1,MGE-IPC2,MGE-IPC3 | IPC |
| MGE-RG1,MGE-RG2,oRG,RG-div1,RG-div2,RG-early,tRG,vRG | RG |
| Microglia | Microglia |
| Endothelial | Endothelial |
| Mural | Mural |
| OPC | OPC |
| **Trevino et al. 2021, Dataset 4** | |
| CGEIN,MGEIN | IN |
| earlyRG,lateRG,tRG | RG |
| EC | Endothelial |
| GluN1,GluN2,GluN3,GluN4,GluN5,GluN6,GluN7,GluN8 | EN |
| MG | Microglia |
| OPC_Oligo | OPC |
| Peric | Mural |
| **Bhaduri et al. 2020 (brain), Dataset 9** | |
| Endothelial | Endothelial |
| ExcitatoryNeuron | EN |
| InhibitoryNeuron | IN |
| IPC | IPC |
| RadialGlia | RG |
| Outlier,Red blood cells | Unknown |
| Mural | Mural |
| Microglia | Microglia |

### Supplementary Figures

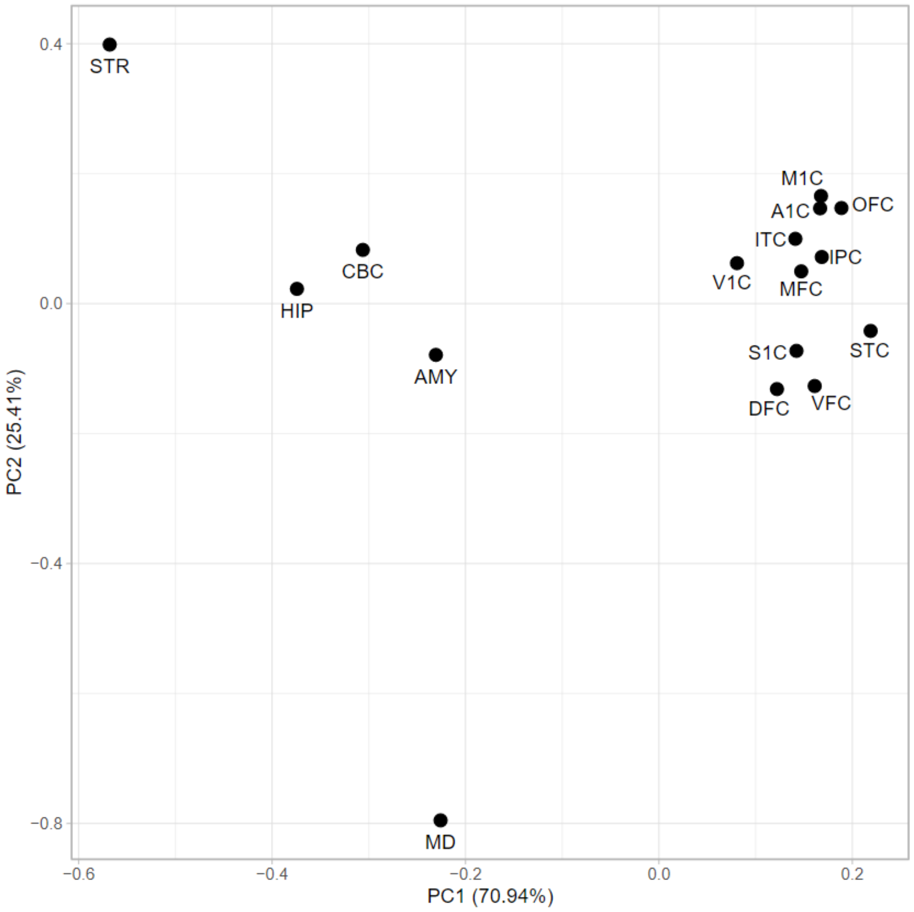


**Figure S1. PCA analysis of brain regions aligned with organoids.** S1C: primary somatosensory(S1) cortex; M1C: primary motor(M1) cortex; OFC: orbital prefrontal cortex; DFC: dorsolateral prefrontal cortex; MFC: medial prefrontal cortex; VFC: ventrolateral prefrontal cortex; STR: stratum; HIP: hippocampus; AMY: amygdala; IPC: posterior inferior parietal cortex; A1C: primary auditory(A1) cortex; V1C: primary visual(V1) cortex; STC: superior temporal cortex; ITC: inferior temporal cortex; MD: mediodorsal nucleus of the thalamus; CBC: cerebella cortex.

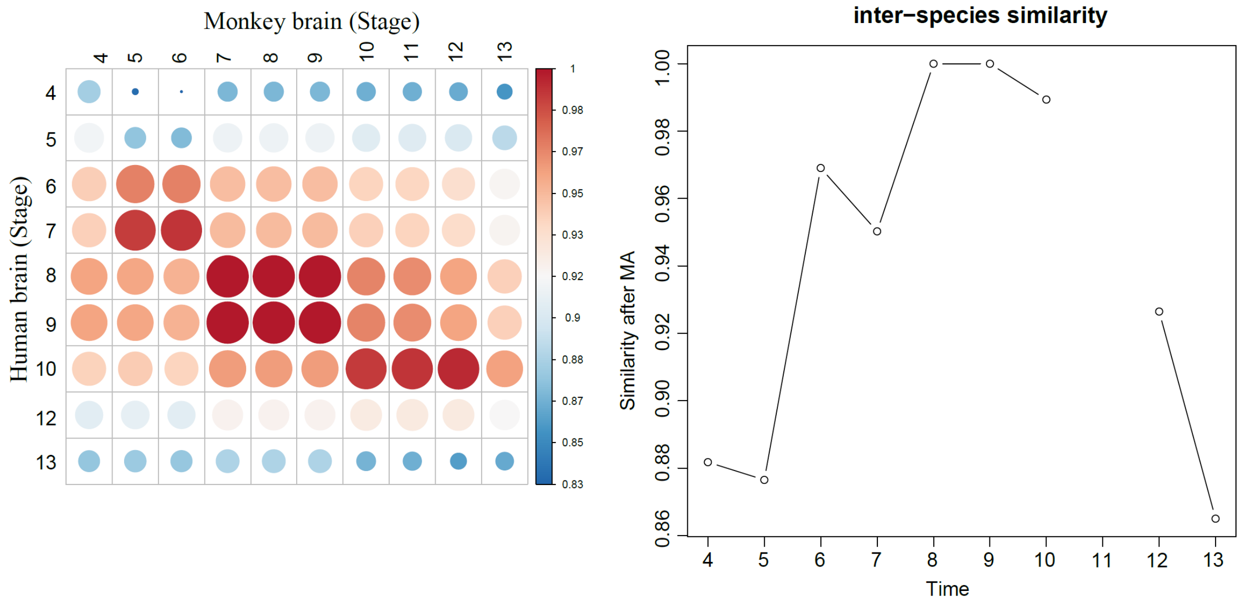


**Figure S2. Human brain VERSUS monkey brain.** The predicted developmental time (Zhu Y. et.al., 2018) for monkey brains was used for the alignment.

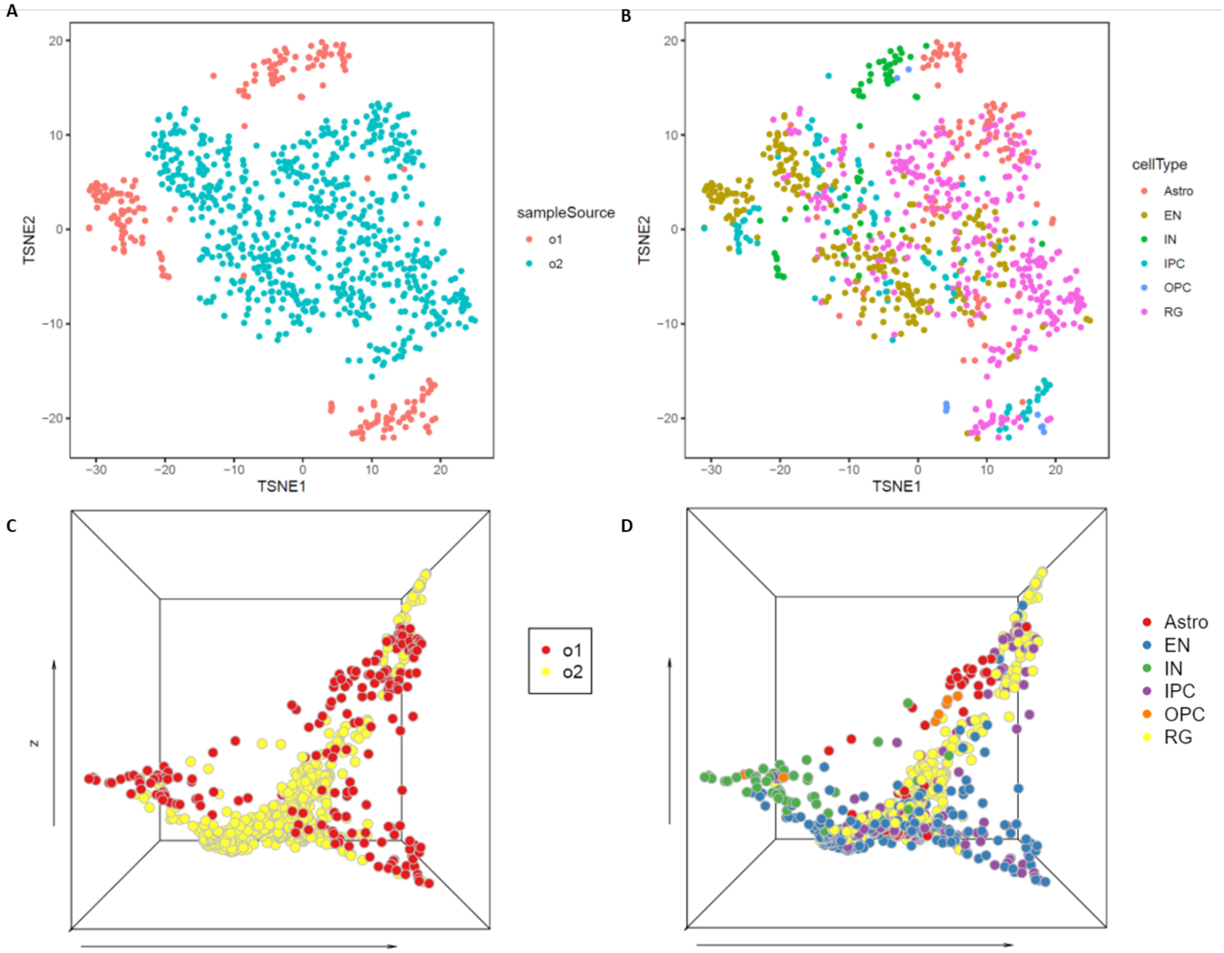


**Figure S3. Minimum batch effects across merged organoid samples were observed by tSNE and BOMA.** A). tSNE analysis of pseudo-cells colored by the corresponding sample sources dataset. The datasets are: Organoid 1(o1, Birey et al. 2017) and Organoid 2(o2, Bhaduri et al. 2020). B). tSNE analysis of pseudo-cells colored by the corresponding cell types. C). 3D scatterplot of pseudo-cells colored by sample sources after BOMA; D). 3D scatterplot of pseudo-cells colored by cell types after BOMA.

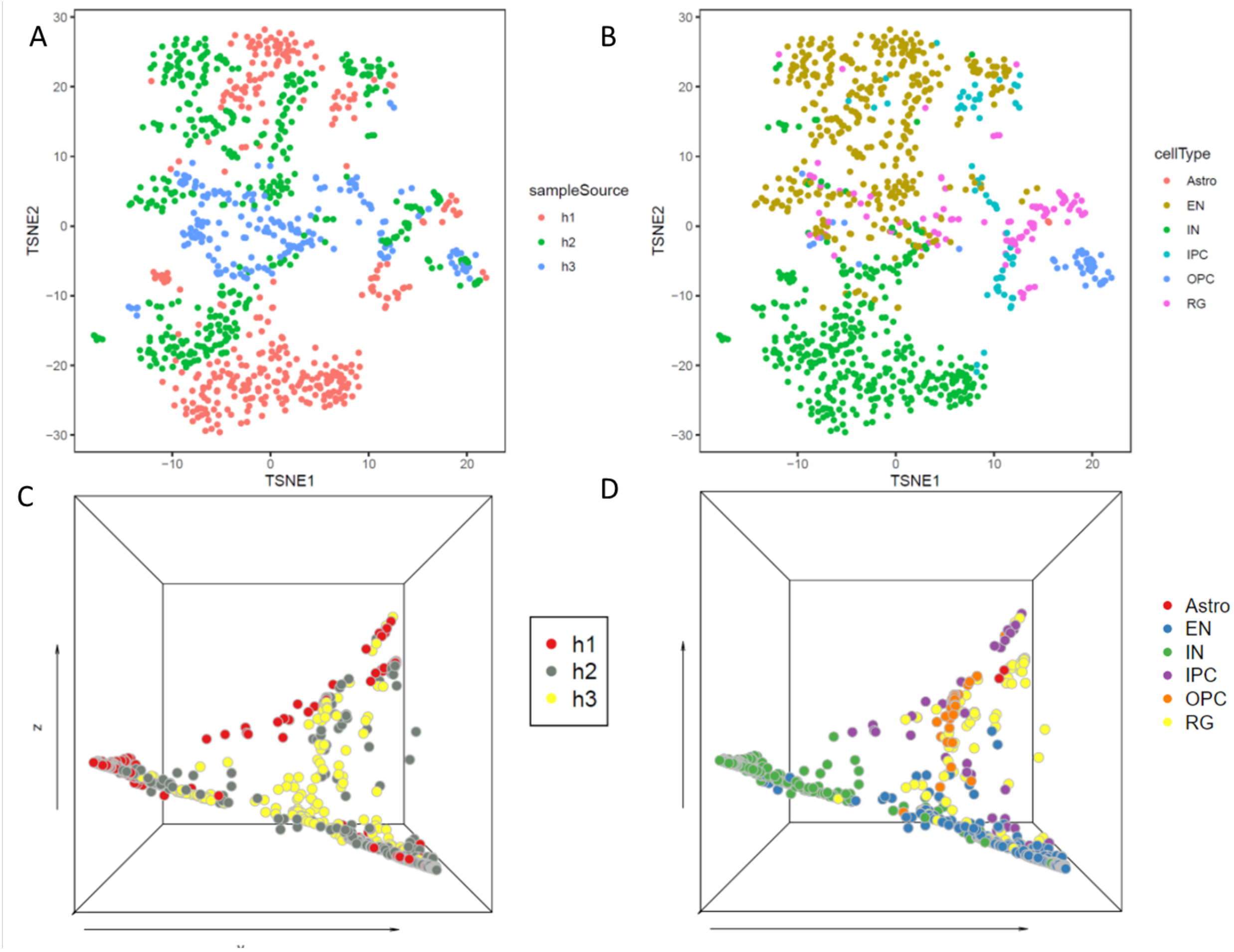


**Figure S4. Minimum batch effects across merged brain samples were observed by tSNE and BOMA.** A). tSNE analysis of pseudo-cells colored by the corresponding sample sources dataset. The datasets are: Human-brain 1(h1, Nowakowski 2017), Human-brain 2(h2, Trevino et al. 2021), Human-brain 3(h3, Bhaduri et al. 2020). E-H). B). tSNE analysis of pseudo-cells colored by the corresponding cell types. C). 3D scatterplot of pseudo-cells colored by sample sources after BOMA; D). 3D scatterplot of pseudo-cells colored by cell types after BOMA.

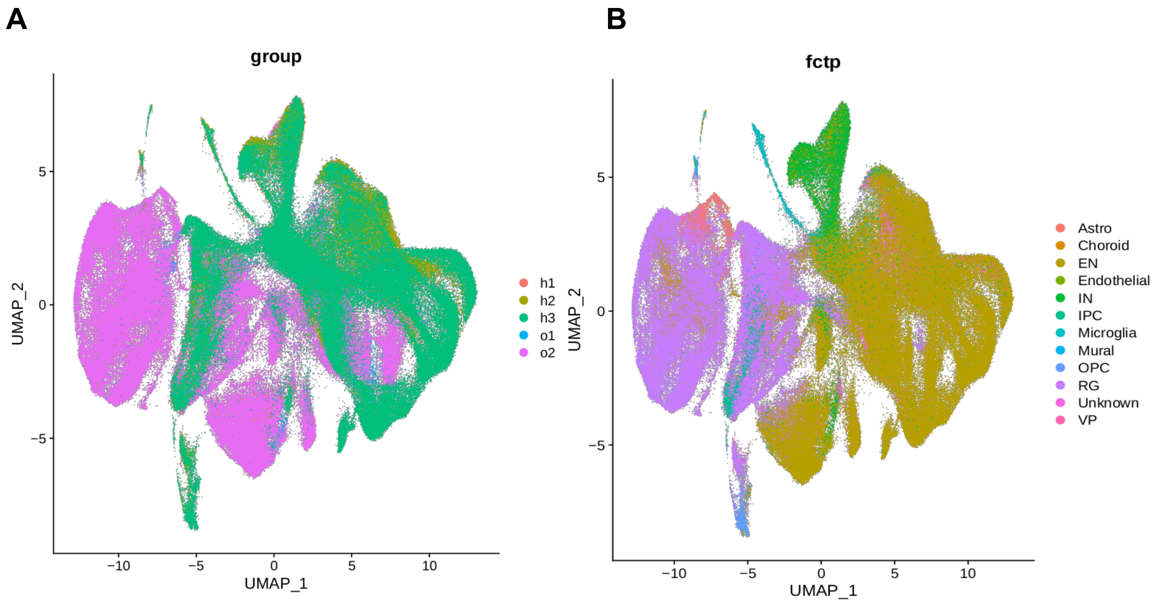


**Figure S5. Human brain and organoid single-cells integrated by Seurat.** A).UMAP colored by datasets. The datasets are h1(human1, Nowakowski 2017), h2(human2, Trevino et al. 2021), h3(human3, Bhaduri et al. 2020), o1(organoid1, Birey et al. 2017) and o2(organoid2, Bhaduri et al. 2020). B). UMAP colored by cell-types

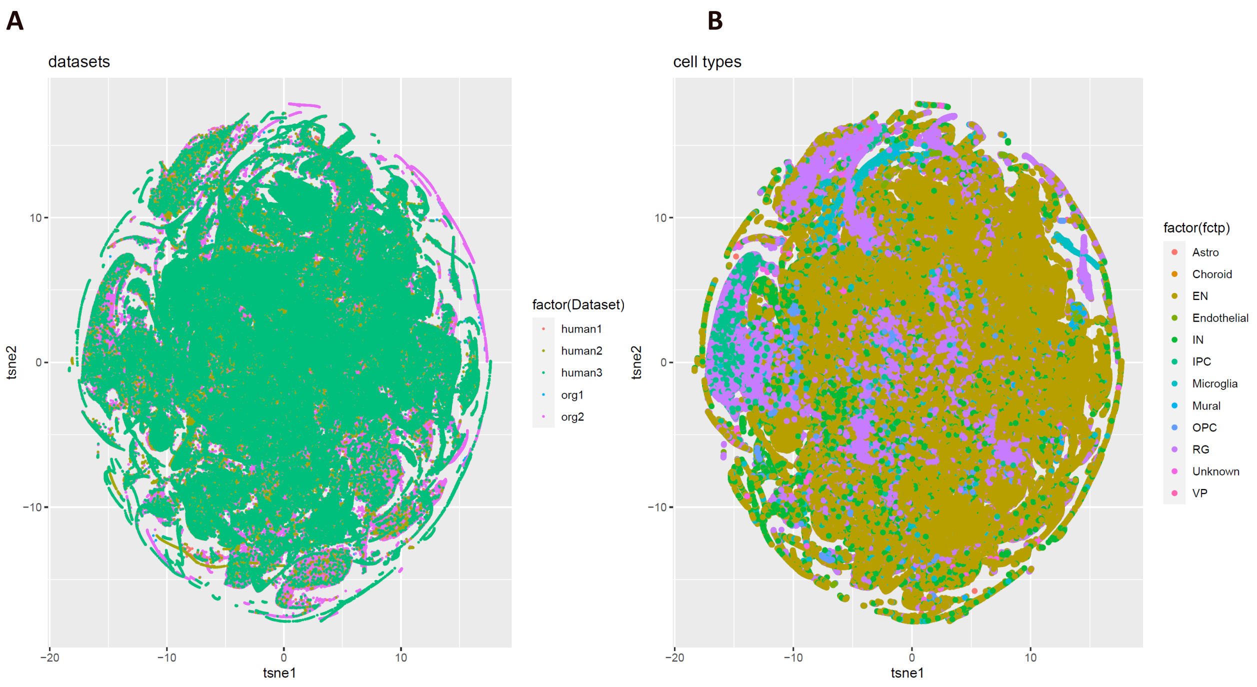


**Figure S6. Human brain and organoid single-cells integrated by Liger.** Tsne colored by datasets. The datasets are human1(Nowakowski 2017), human2(Trevino et al. 2021), human3(Bhaduri et al. 2020), org1(organoid1, Birey et al. 2017) and org2(organoid2, Bhaduri et al. 2020). B). Tsne colored by cell-types

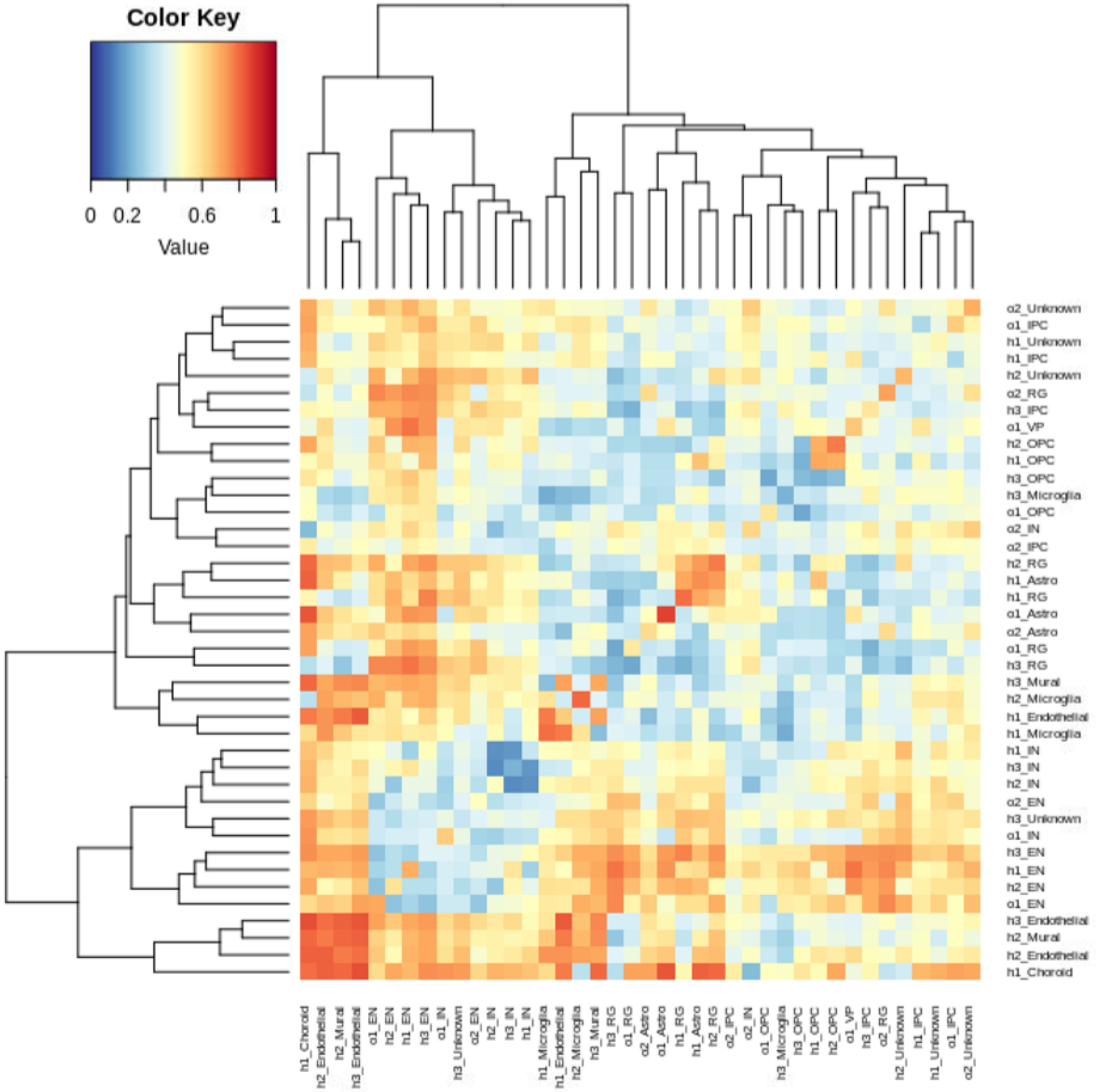


**Figure S7. Cell-types similarity across scRNA-seq datsets quantified by MetaNeighbor.** Heatmap generated by applying MetaNeighbor on 10% randomly sampled single-cells from integrated scRNA-seq dataset in Figure 5. The datasets are h1(human1, Nowakowski 2017), h2(human2, Trevino et al. 2021), h3(human3, Bhaduri et al. 2020), o1(organoid1, Birey et al. 2017) and o2(organoid2, Bhaduri et al. 2020)

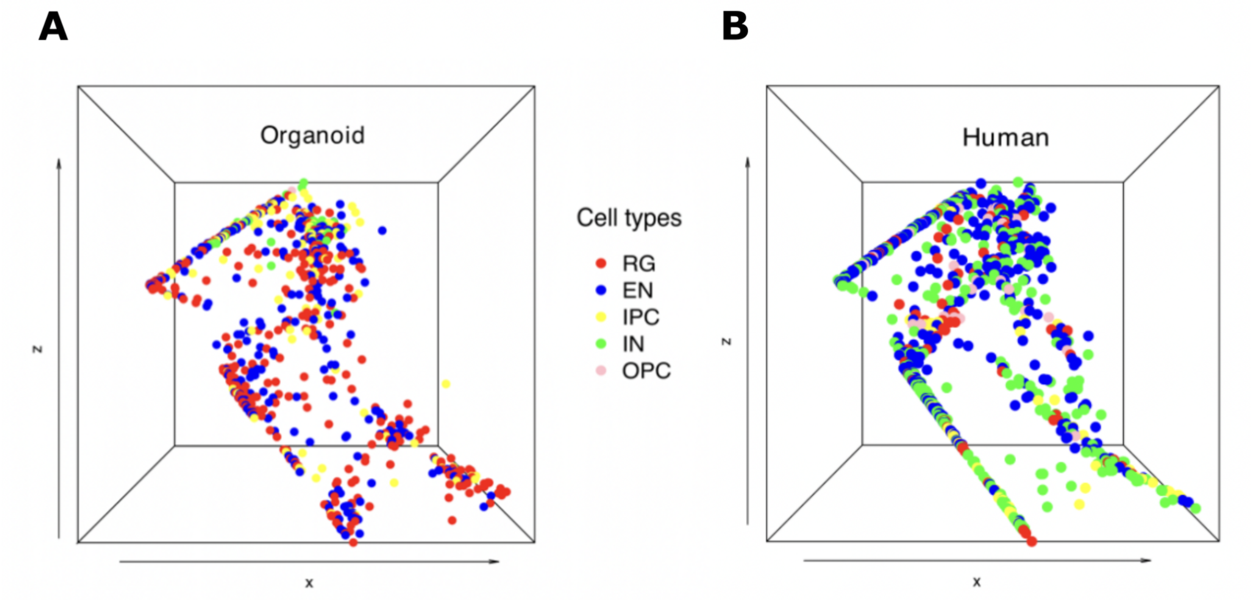


**Figure S8. Human brain and organoid pseudo-cells aligned by Unioncom.** A). The 3D scatter plot shows the alignment results for 766 organoid pseudo-cells. B). The 3D scatter plot shows the alignment results for 1,016 human brain pseudo-cells. The dots are colored by the cell types. RG cells are marked in red, EN cells are marked in blue, IPC cells are marked in yellow, IN cells are marked in green and OPC cells are marked in pink.

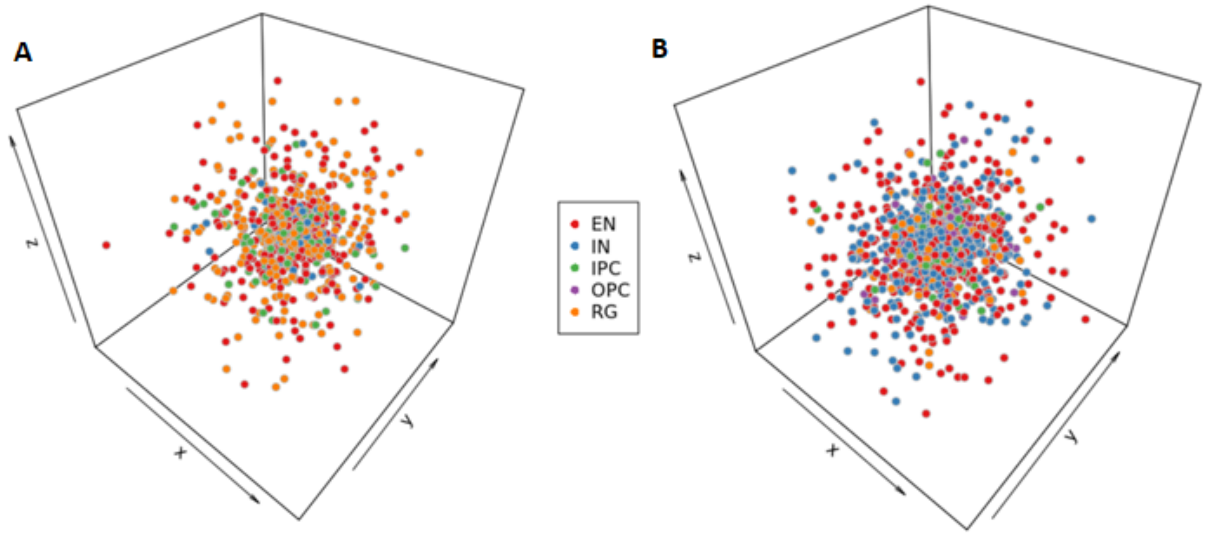


**Figure S9. Human brain and organoid pseudo-cells aligned by MMD-MA.** A). The 3D scatter plot shows the alignment results for 766 organoid pseudo-cells. B). The 3D scatter plot shows the alignment results for 1,016 human brain pseudo-cells. The dots are colored by the cell types.

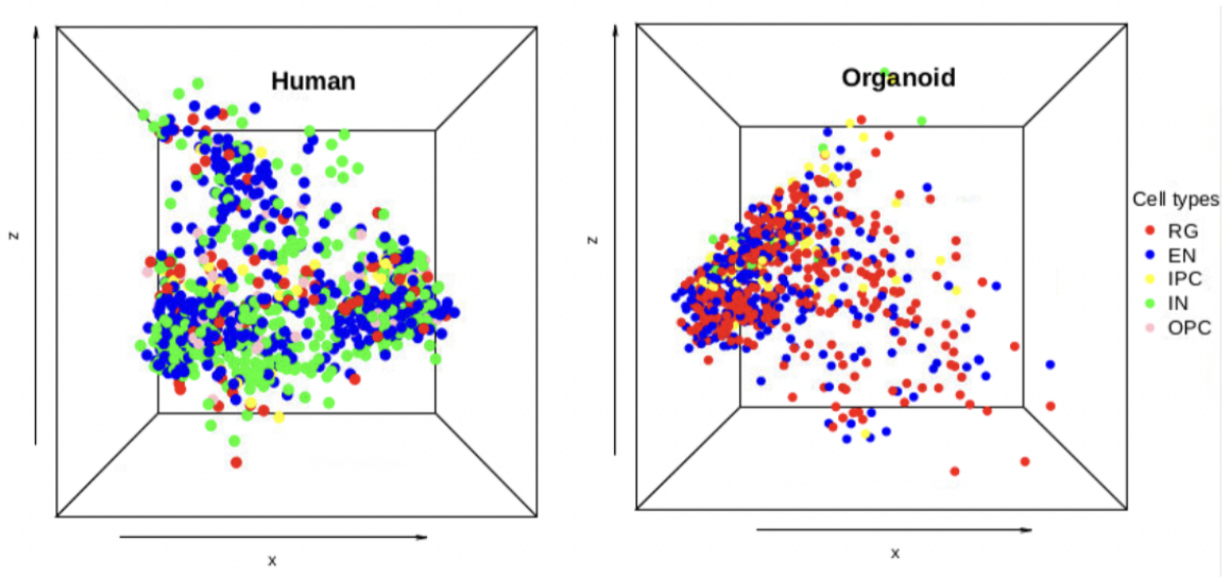


**Figure S10. Human brain and organoid pseudo-cells aligned by SCOT.** A). The 3D scatter plot shows the alignment results for 766 organoid pseudo-cells. B). The 3D scatter plot shows the alignment results for 1,016 human brain pseudo-cells. The dots are colored by the cell types.

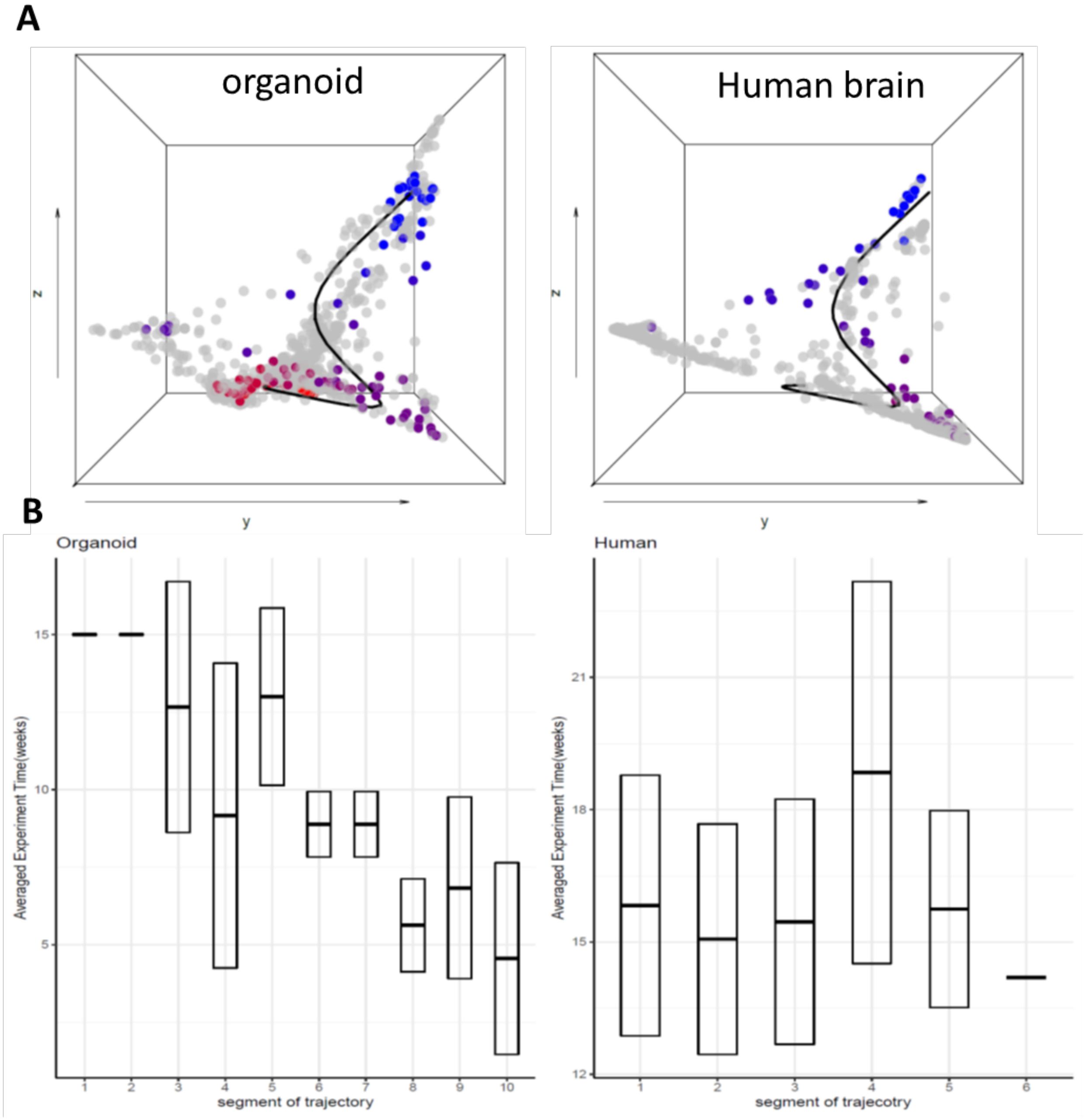


**Figure S11. Trajectory analysis for IPCs.** A). 3D scatter plot shows the inferred trajectory of IPC from the alignment. Organoid (left panel) and brain(right panel) cells are colored by pseudo-time along the trajectory. Blue means earlier while red means later. B). The trajectory was divided into 10 segments and correlate with experiment timepoints. The box plot shows the distribution of experimental time-points.

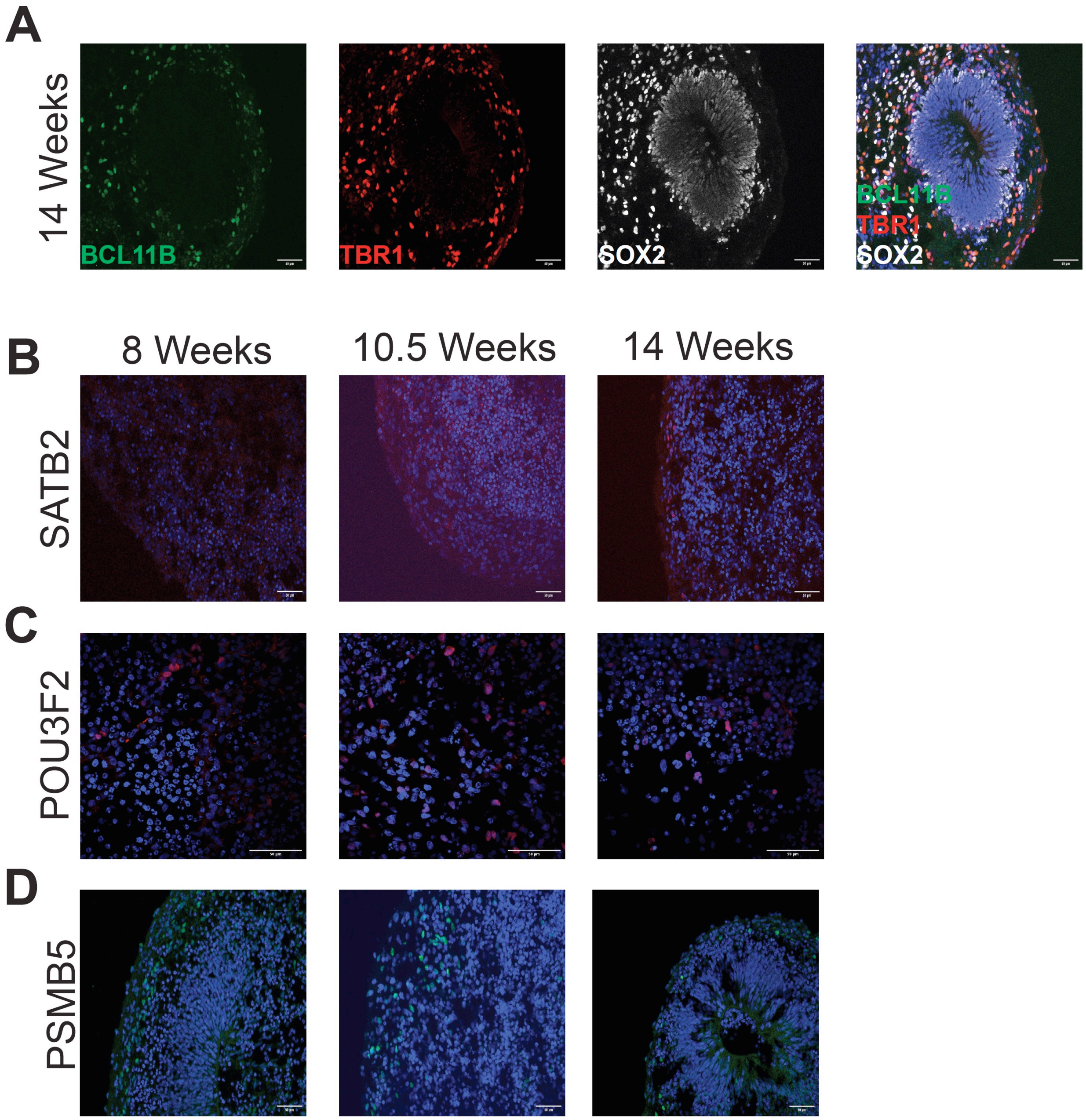


**Figure S12. Representative images of human cortical organoids immunostaining.** (A). Image shows expressing progenitor cells positive for SOX2, deep layer cortical neurons positive for BCL11B and TBR1. Immunostaining of different stages of differentiation in cortical organoids positive for SATB2 (B), POU3F2 (C), and PSMB5 (D) cells. Quantification is shown in Figures 6C, 6F, and 6I. Scale bars, 50μm.

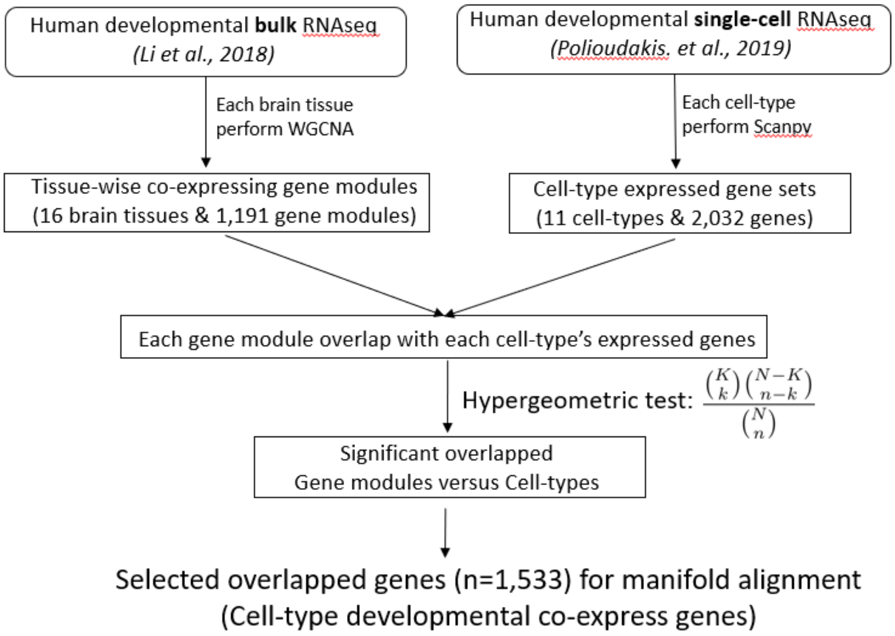


**Figure S13. Flowchart of selecting human brain developmental genes.** BH adjusted P-value <0.01 was used as statistical significance threshold.

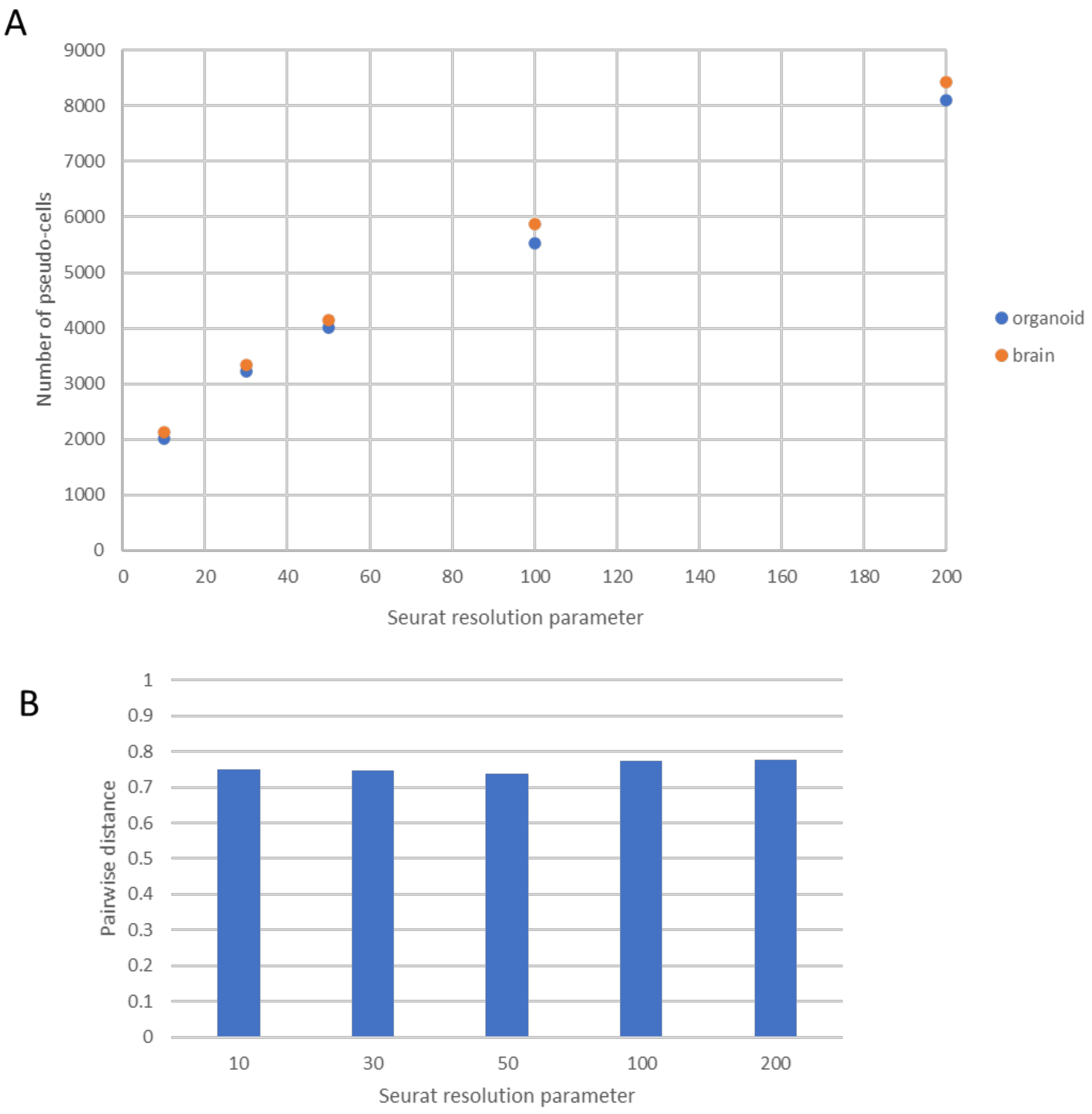


**Figure S14. BOMA is scalable to the input pseudo-cells.** A). Number of pseudo-cells generated at different Seurat resolution; B). Averaged pairwise distances between pseudo-cells of the same cell-type.
